# MapA, a second large RTX adhesin, contributes to biofilm formation by *Pseudomonas fluorescens*

**DOI:** 10.1101/2020.05.11.089839

**Authors:** Alan J. Collins, Alexander B. Pastora, T. Jarrod Smith, Kurt Dahlstrom, George A. O’Toole

## Abstract

Mechanisms by which cells attach to a surface and form a biofilm are diverse and differ greatly between organisms. The Gram-negative, Gammaproteobacterium *Pseudomonas* fluorescens attaches to a surface through the localization of the large type 1-secreted RTX adhesin LapA to the outer surface of the cell. LapA localization to the cell surface is controlled by the activities of a periplasmic protease, LapG and an inner-membrane spanning cyclic di-GMP responsive effector protein, LapD. A previous study identified a second, LapA-like protein encoded in the *P. fluorescens* Pf0-1 genome: Pfl01_1463. However, deletion of this gene had no discernible phenotype under our standard laboratory growth conditions. Here, we identified specific growth conditions wherein, Pfl01_1463, hereafter called MapA (<u>M</u>edium <u>A</u>dhesion <u>P</u>rotein A) is a functional adhesin contributing to biofilm formation. This adhesin, like LapA, appears to be secreted through a Lap-related type 1 secretion machinery. We show MapA involvement in biofilm formation is also controlled by LapD and LapG, and that the differing roles of LapA and MapA in biofilm formation are achieved, at least in part, through the differences in the sequences of the two adhesins and their differential, cyclic di-GMP-dependent transcriptional regulation. This differential regulation appears to lead to different distributions of the expression of *lapA* and *mapA* within a biofilm. Our data indicate that the mechanisms by which a cell forms a biofilm and the components of a biofilm matrix can differ depending on growth conditions in the biofilm.

**Importance:** Adhesins are critical for the formation and maturation of bacterial biofilms. We identify a second adhesin in *P. fluorescens*, called MapA, which appears to play a role in biofilm maturation and whose regulation is distinct from the previously reported LapA adhesin, which is critical for biofilm initiation. Analysis of bacterial adhesins show that LapA-like and MapA-like adhesins are found broadly in Pseudomonads and related organisms, indicating that the utilization of different suites of adhesins may be broadly important in the Gammaproteobacteria.

## Introduction

Biofilm formation is a critical mode of growth for many microorganisms, including bacteria (1), archaea (2) and fungi (3). The mechanisms by which individual cells can attach to abiotic surfaces, and to each other, to form a biofilm are highly varied. However, in all cases formation of a biofilm involves the production of adhesive molecules by the cells. These molecules are most commonly polysaccharide- or protein-based, and these factors remain adhered to the substratum while also often remaining tethered to the cells that produced them (4–8).

In bacteria many different mechanisms of surface attachment by cells have been described. The biofilm matrix is typically composed of a variety of molecules, including a combination of polysaccharides, proteins, and extracellular DNA used to join cells together into a biofilm (4, 9–12). In general a combination of multiple molecules make up the biofilm matrix, but some organisms use different molecules as their major biofilm component. For example, the production of a polysaccharide called *Vibrio* polysaccharide (VPS) and the matrix proteins RbmA, RbmC and Bap1 by *Vibrio cholerae* (13–15) or the production of amyloid proteins called curli fibers by *Escherichia coli* (16). For many bacteria, biofilm formation is important for the effective colonization of their natural niches in the soil, on plants, or in the water column (17–19). Biofilm formation has also been implicated as an important virulence factor for many pathogenic bacteria (20–22). It is therefore important to understand both the mechanisms by which bacteria can form a biofilm, and the ways in which that process is regulated.

An organism with one of the best-understood mechanisms of surface attachment and regulation of biofilm formation is *Pseudomonas fluorescens*. In *P. fluorescens*, surface attachment is predominantly achieved through the activity of a single large adhesion protein, LapA (23, 24). LapA is a non-canonical Type 1-secreted, RTX adhesin which is composed of a C-terminal secretion signal, as well as a central portion containing numerous repeat domains, multiple calcium binding domains, and a von Willibrand A domain that all contribute to adhesion (25–27). LapA also has a N-terminal globular domain that folds while the protein is in the process of being secreted thus anchoring the protein in its TolC-like outer-membrane pore (28, 29). LapA is a ∼520 kDa adhesin (25–27), and was thus named the <u>L</u>arge <u>A</u>dhesion <u>P</u>rotein A, and for *lapa*, the Spanish word for limpet, a marine mollusk that attaches tightly to rocks and other surfaces (23). The localization of LapA to the cell surface is dependent on the activity of two regulatory proteins: LapG, a periplasmic cysteine protease that can proteolytically process LapA, removing the globular N-terminus and releasing LapA from the cell surface; and LapD, an inner-membrane localized GGDEF/EAL-containing cyclic di-GMP-responsive effector protein which binds LapG in response to cytoplasmic cyclic di-GMP, preventing its access to the N-terminus of LapA (25, 28, 30–34). Together, LapD and LapG function to transduce cytoplasmic cyclic di-GMP signals across the inner-membrane and determine whether LapA is retained at the cell surface or not (28, 34, 35). While the biofilms formed by some lab strains and environmental isolates of *P. fluorescens* have been shown to contain appreciable levels of polysaccharides (36, 37), in the best studied lab strain, Pf0-1, *P. fluorescens* polysaccharides have not previously been shown to play an appreciable role in biofilm formation. Instead, LapA has been shown to be the key determinant of biofilm formation (23, 28, 31). The simplicity of the use of a single adhesin protein to determine surface attachment and biofilm formation has made *P. fluorescens* an attractive model system for the study of biofilm formation and its regulation by c-di-GMP.

In this study we demonstrate that biofilm formation in *P. fluorescens* is not determined solely by the action of LapA, but instead the action of two, related proteins: LapA and the herein described <u>M</u>edium <u>A</u>dhesion <u>P</u>rotein A (MapA). We show that, while these two proteins share much of the regulatory machinery controlling cell-surface localization, the differences in transcriptional regulation, their differential expression within mature biofilms, and their distinct sizes and sequences suggest that these two large adhesins play different roles in the development of a mature biofilm by *P. fluorescens*. We also demonstrate that LapA- and MapA-like proteins can be found in various combinations in Gammaproteobacteria.

## Results

### LapA is not necessary for the formation of a biofilm under all media conditions

Previous work characterizing the role of the adhesive protein LapA in biofilm formation had suggested that biofilm formation by *P. fluorescens* Pf0-1 is dependent on the function of LapA (23, 25, 28, 29). However, recent work by Smith *et al*. (29) indicated that *P. fluorescens* Pf0-1 encodes a second putative large adhesin. LapA is approximately 520 kDa, while this second adhesin, encoded by the Pfl01_1463 gene, is predicted to be approximately 300 kDa. We therefore named this new putative adhesin MapA (<u>M</u>edium <u>A</u>dhesion <u>P</u>rotein A) as the data shown below supports a role for MapA as a biofilm adhesin.

We first assessed biofilm formation by these mutants in the medium K10-T, a high-phosphate medium containing glycerol and tryptone as carbon and nitrogen sources that is commonly used in studies of *P. fluorescens* Pf0-1 biofilm formation. After 6 hours of growth in this medium, the Δ*lapA* mutant formed no biofilm, similarly to the Δ*lapA*Δ*mapA* double mutant (Figure 1A). This pattern was also observed after 16 hours of growth in K10-T medium (Figure 1B, E).

**Figure 1.**
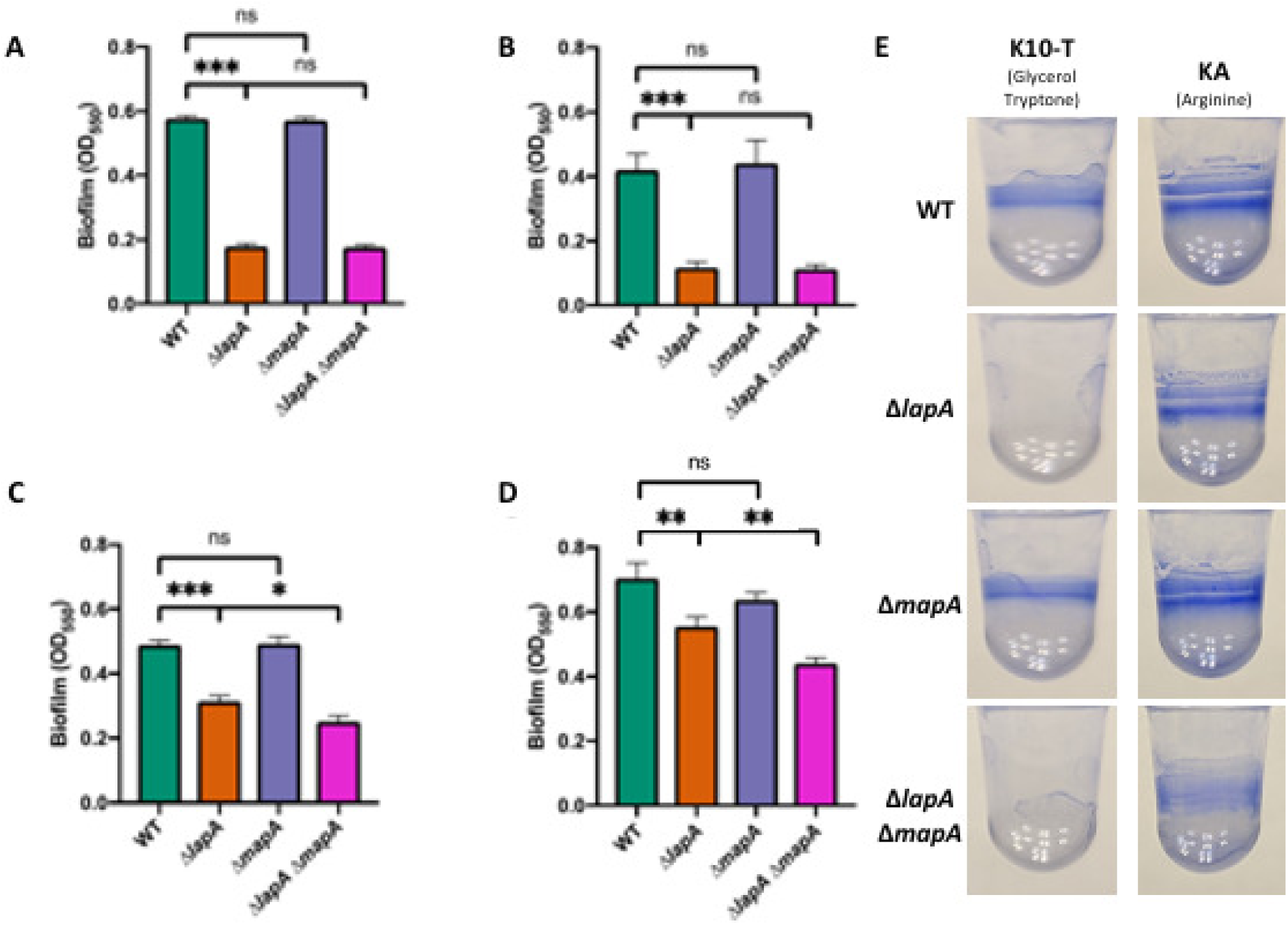
Biofilm formation by adhesin mutants. (A-D) Quantification of biofilm formed by WT *P. fluorescens* Pf0-1, Δ*lapA*, Δ*mapA*, and the Δ*lapA*Δ*mapA* double mutant after (A) 6 hours of growth in K10-T medium, (B) 16 hours of growth in K10-T medium, (C) 6 hours of growth in KA medium, and (D) 16 hours of growth in KA medium. Statistical tests: hypothesis testing was conducted in Graphpad Prism 8 using two tailed t tests. P values are indicated by asterisks as follows: * P < 0.05, ** P < 0.01, *** P < 0.001, ns P > 0.05. (E) Representative image of biofilm formed WT *P. fluorescens* Pf0-1, Δ*lapA*, Δ*mapA*, and the Δ*lapA*Δ*mapA* mutants after 16 hours of growth in the indicated medium and stained with 0.1% (w/v) crystal violet.

In order to find a role for MapA in biofilm formation, we sought to assess the biofilm formed by a Δ*lapA*, or Δ*mapA* single mutant, or a Δ*lapA*Δ*mapA* double mutant under a growth condition that fosters robust biofilms. Arginine has been shown to strongly promote biofilm formation by *Pseudomonas* species, likely through stimulating the production of c-di-GMP (38–41). In addition, arginine is commonly found in root exudates and is therefore likely encountered by *P. fluorescens* when growing in the soil (42–44). After trying several medium formulations (data not shown), we found that the condition in which a Δ*lapA* mutant is most capable of biofilm formation is a modification of K10-T medium wherein no glycerol or tryptone is added, but rather it is supplemented with 0.4% (w/v) L-Arginine HCl (henceforth referred to as KA medium), as described in the Materials and Methods.

When grown in KA medium, WT *P. fluorescens* Pf0-1 can form a biofilm by 6 hours, although less than the biofilm typically observed in K10-T medium (Figure 1C). After 6 hours of growth, the Δ*lapA* mutant has a significantly higher level of biomass attached to the well than the Δ*lapA*Δ*mapA* mutant, although the single Δ*mapA* shows no reduction in biofilm compared to the WT. When cells were allowed to grow for 16 hours in KA medium the relative deficit in biofilm formation by the Δ*lapA* mutant was reduced, while the significant deficit of the Δ*lapA*Δ*mapA* mutant was still observed (Figure 1D, E). In addition, the biofilm formed by the Δ*mapA* mutant in KA medium after 16 hours of growth was slightly (but not significantly) lower than that of WT (P=0.09, unpaired two-tailed t test). Interestingly, when the Δ*lapA*Δ*mapA* mutant was grown in KA for 16 hours, there was still modest diffuse staining observed in the well (Figure 1E), perhaps indicating an additional factor contributing to biofilm formation under these conditions. These data indicate that MapA is capable of significantly contributing to biofilm in some medium conditions and is able to do so in the absence of LapA when grown in KA medium.

### Genetic studies indicate that MapA localization is controlled by the regulatory proteins LapD and LapG

Localization of LapA to the cell surface is dependent on the activity of the proteins LapD and LapG. The structure of the N-terminal portion of LapA is essential for its retention in the outer membrane pore LapE and thus for anchoring LapA to the bacterial cell surface (25, 29). *In silico* analysis of the N-terminal portion of LapA and MapA indicates a high degree of sequence and secondary structure similarity as well as presence of the functionally important TAAG motif required for LapG processing (35) (Figure 2A, red box).

**Figure 2.**
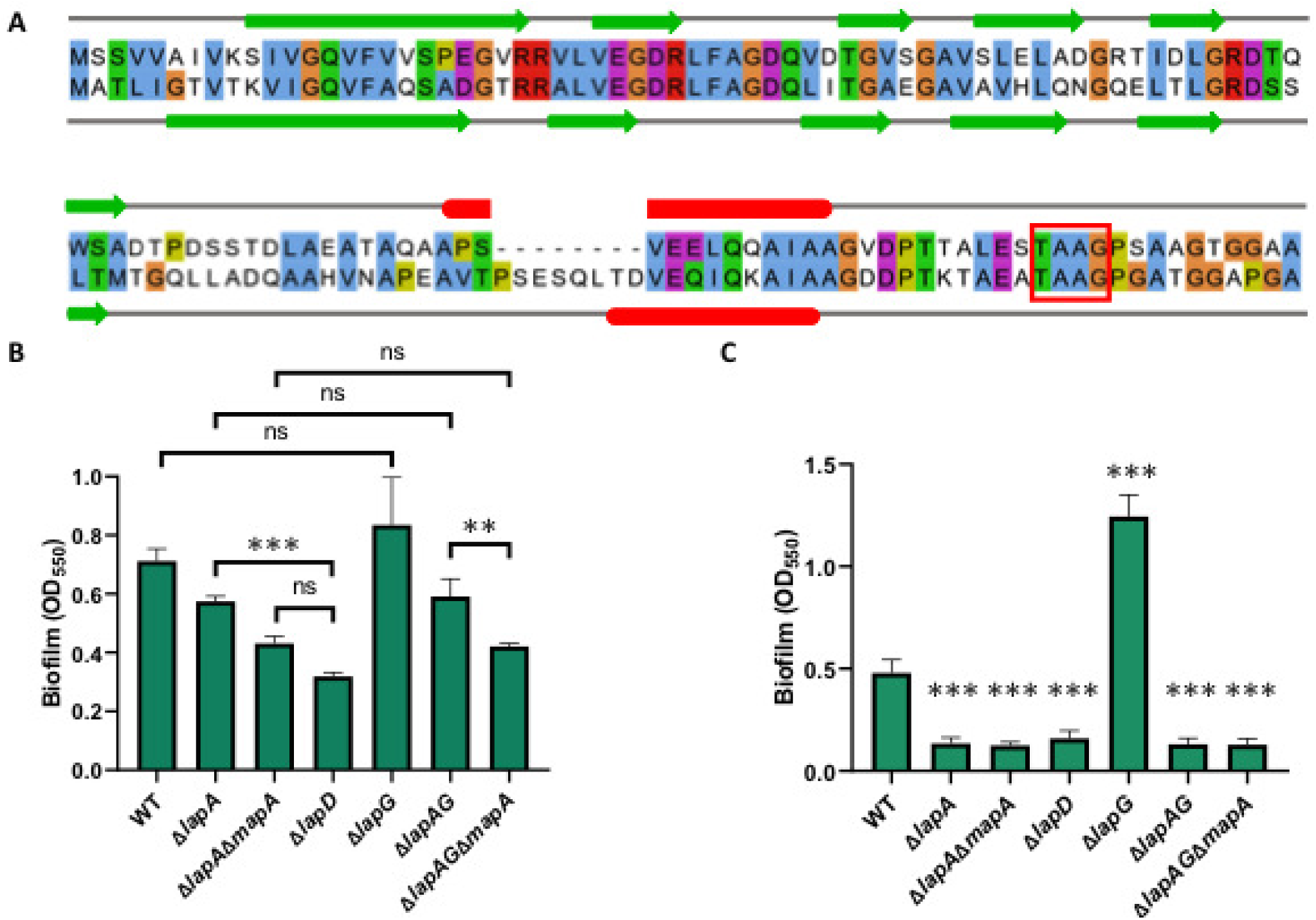
LapD and LapG contribute to MapA-dependent biofilm formation. (A) Alignment of LapA and MapA N-terminal portions generated using MUSCLE (74) and visualized using Jalview (93) with secondary structure prediction using JPred4 (94). Similar residues are colored. Predicted secondary structure is indicated by colored shapes: Green arrows = beta sheets, Red blocks = alpha helix. (B) Quantification of biofilm assay assessing biofilm formation of adhesin and regulatory mutants grown for 16 hours in KA medium. (C) Quantification of biofilm assay assessing biofilm formation of adhesin and regulatory mutants grown for 16 hours in K10-T medium. Statistical tests: hypothesis testing was conducted in Graphpad Prism 8 using two tailed t tests with Holm-Sidak’s multiple comparison correction. P values are indicated by asterisks as follows: ** P < 0.01, *** P < 0.001, ns P > 0.05. In panel C all asterisks denote P values of t tests between WT and the indicated mutant with Holm-Sidak’s multiple comparison correction.

Previous work demonstrated that the regulatory protease, LapG is capable of proteolytically processing the N-terminal portion of MapA in vitro (29). To assess the regulatory role of LapD and LapG in the localization of MapA we examined the level of biofilm formation of mutants lacking *lapD* or *lapG* when grown in KA medium (Figure 2B). We found that, while a Δ*lapA* mutant can form a biofilm in KA medium, in a Δ*lapD* mutant the biofilm is significantly reduced compared to the Δ*lapA* mutant, consistent with LapD being required to prevent constitutive processing of LapA and MapA by LapG. We also observe a reduction in the biofilm formed by the Δ*lapAG*Δ*mapA* compared to the Δ*lapAG* mutant, indicating that MapA does contribute to the biofilm formed under these conditions. Interestingly, when cells were grown in KA medium (Fig. 2B), no significant difference in biofilm formation was observed between WT and the Δ*lapG* mutant, or the Δ*lapA* and Δ*lapAG* mutant. These data suggest that, while LapG is capable of processing both LapA and MapA, cells grown in KA medium have very little LapG activity so the loss of LapG does not lead to an appreciable increase of biofilm formation, contrary to cells grown in K10-T medium (Figure 2C). Finally, when grown in KA medium, the Δ*lapD* mutant forms less biofilm than the Δ*lapA*Δ*mapA* mutant, although this difference is not statistically significant after multiple comparisons correction. This observation may indicate the existence of an additional, unknown LapD-regulated factor that contributes to biofilm formation under these conditions, an observation consistent with the biofilm data shown in Figure 1D-E (see above). Taken together, these genetic data are consistent with the previous biochemical data (29) indicating that MapA is a substrate of LapG and its localization is likely regulated by LapD.

### MapA is localized to the cell surface

The sequence similarity between the N-terminus of LapA and MapA, the role for MapA in biofilm formation, the impact of loss of function in LapD/LapG on MapA-dependent biofilm, as shown here, and the published observation that the N-terminus of MapA is processed by LapG (29) are all consistent with the conclusion that MapA is a cell-surface localized adhesin. To assess the cell surface localization of MapA, we built an HA-tagged variant of MapA – this strain showed no difference in biofilm formation compared to the untagged WT strain when grown in KA medium (Fig. 3A, left); we similarly saw no impact of the HA-tagging of MapA on the biofilm formed in the *lapA* mutant background (Fig. 3A, right).

**Figure 3.**
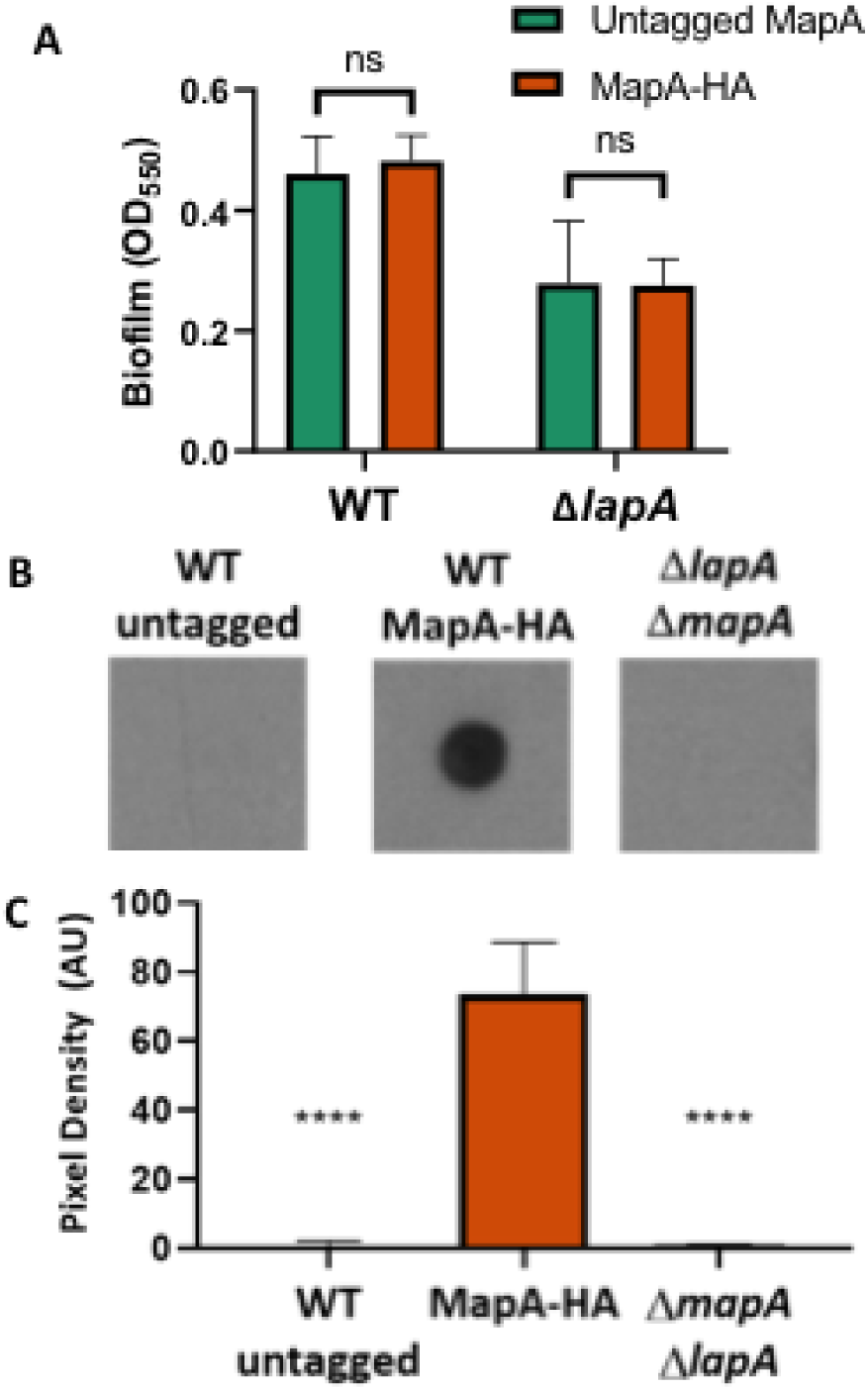
MapA is localized to the cell surface. (A) Quantification of crystal violet staining of wells of 96-well plate after 16 hours of growth in KA. *P. fluorescens* Pf0-1 in both the untagged MapA and the tagged MapA variant bearing 3 tandem HA tags. The phenotype is shown either in strains carrying a functional LapA or a strain for which the *lapA* gene has been deleted. When compared using student’s t-test neither the *P. fluorescens* Pf0-1 (p =0.25) nor the Δ*lapA* mutant (p =0.9) formed a significantly different level of biofilm when bearing an HA-tagged variant of MapA. (B) Shown is a representative dot blot of cell-surface localized MapA for the indicated strains. The WT (untagged strains) and a strain deleted for both the *mapA* and *lapA* gene serve as negative controls. (C) Three independent dot blot experiments were quantified using ImageJ as follows. The pixel values for the blots were inverted for ease of plotting. The region of interest (ROI) was defined for the control LapA dot blot (not shown here), then the mean gray value was determined for each spot using the pre-defined ROI. The background was subtracted from each spot; the background was determined by measuring the mean gray value in a section of the blot not containing any samples. The adjusted mean pixel density was plotted here for three biological replicates. There was a significant reduction in signal for both control strains compared to the strain expressing MapA-HA as assessed by t test, * P < 0.05, *** P < 0.001, ns P > 0.05.

We next performed dot blot studies that specifically assess the MapA associated with the bacterial cell surface. In these studies, cells are harvested by centrifugation, washed, spotted on a nitrocellulose filter, then probed with an antibody to the HA-tagged variant to MapA. As shown in Figure 3B-C, while no signal is detected for the untagged WT strain (left), a strong signal is detected for the strain carrying the HA-tagged MapA (center), and as an additional negative control, this signal is absent from a strain in which both the *lapA* and *mapA* genes are deleted (right). These data are consistent with MapA, like LapA, being cell-surface localized.

### Genetic evidence that components of the T1SS of LapA may be promiscuous

Homologs of LapA are commonly encoded adjacent to their cognate T1SS (23, 45). This is also true for MapA. The *mapA* gene is adjacent to three genes encoding a putative T1SS (Figure 4). We have named these genes *mapB, mapC*, and *mapE*, encoding a predicted inner membrane-spanning ABC transporter, a membrane fusion protein, and a outer membrane pore protein, respectively. Given that TolC, a homolog of LapE and MapE, is capable of functioning in association with multiple different ABC transporters and membrane fusion proteins (46–50) and the sequence similarity between these proteins (both LapE and MapE share roughly 35% similarity with TolC), we next sought to assess the extent to which LapA and MapA might share the components of their secretion systems. To investigate the ability of LapA and MapA to be secreted through either secretion machinery, we assessed the level of biofilm formed by mutants lacking combinations of adhesin and secretion machinery genes grown for 16 hours in KA.

**Figure 4.**
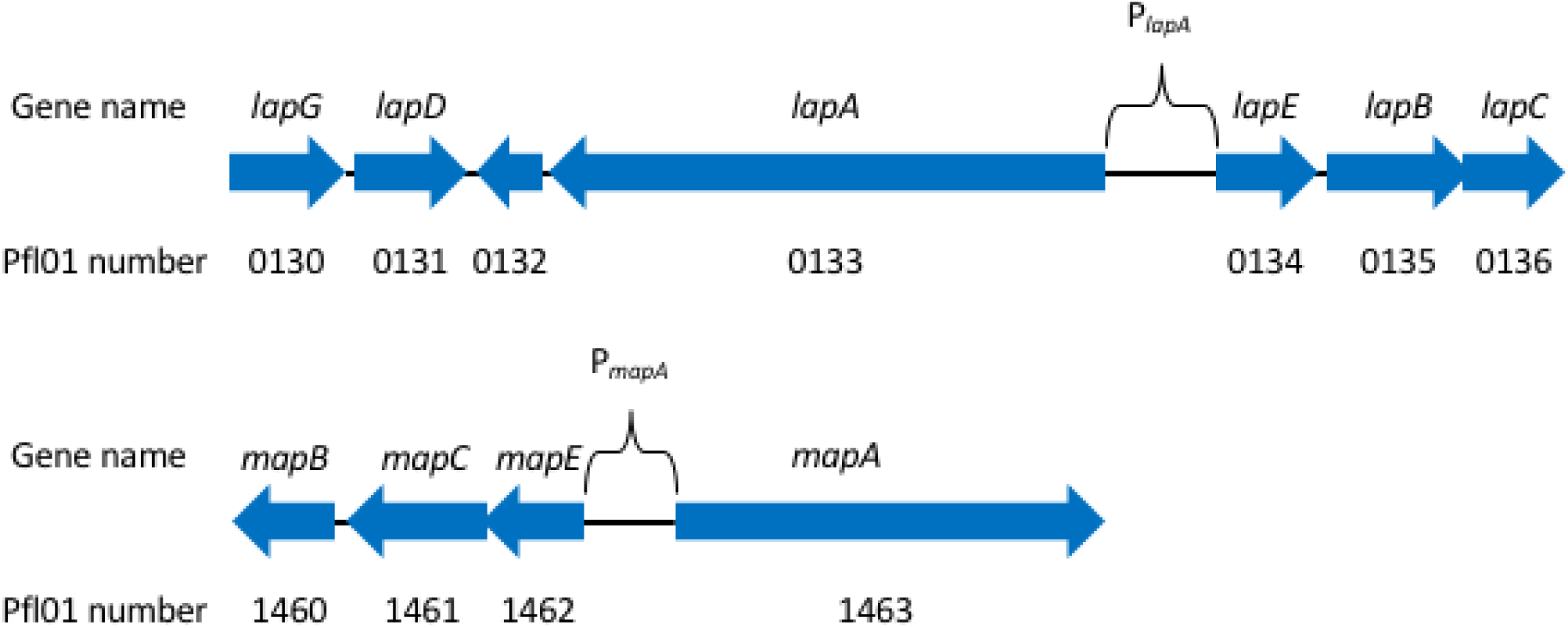
Schematic of the *lap* and *map* loci. The organization of the genes encoding the LapA and MapA adhesins and their secretion machinery are shown. Gene names are indicated above the arrows and the *P. fluorescens* Pfl01 gene numbers are indicated below the arrows. Arrows indicate the location and direction of transcription of the ORFs. Arrow direction indicates which orientation the gene is encoded. Not to scale.

The biofilms produced by the Δ*lapA*, Δ*mapA* and Δ*mapE* mutants were not different from one another but were significantly reduced compared to WT (Figure 5A). The biofilm formed by the Δ*lapA*Δ*mapE* mutant was reduced compared to both the Δ*lapA* and the Δ*mapE* mutants but was not as low as that produced by the Δ*lapA*Δ*mapA* or the Δ*lapE*Δ*mapE* mutants. This intermediate biofilm indicates that when LapA and MapE are absent, MapA is likely able to be secreted, at least in part, via some portion of the LapBCE T1SS and thereby contribute to biofilm formation.

**Figure 5.**
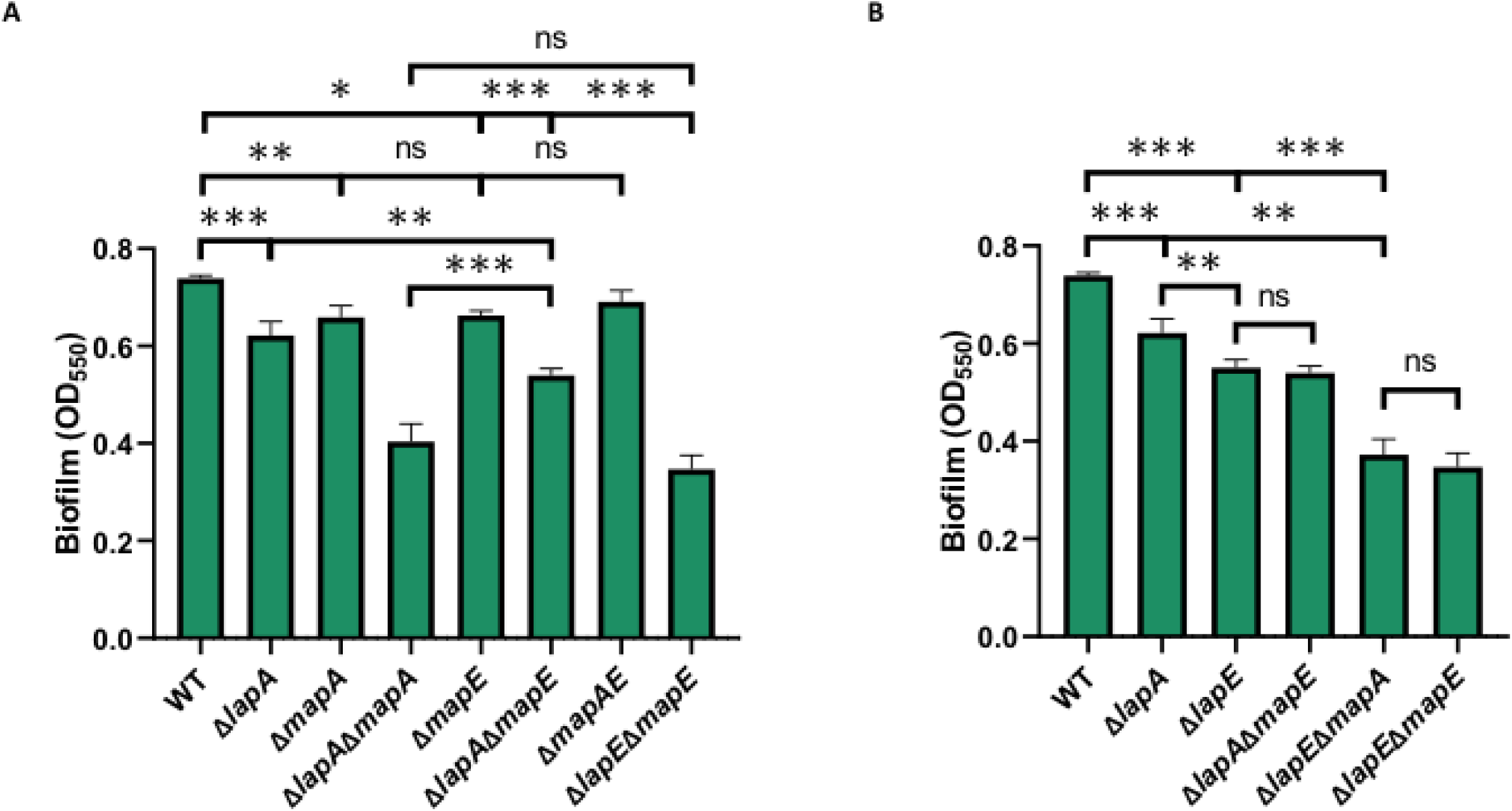
Genetic evidence that MapA can utilize the Lap secretion system. (A) Biofilm formation of strains in KA medium carrying mutations in adhesin genes and genes encoding the Map secretion system outer membrane pore component. (B) Biofilm formation in KA medium for strains carrying mutations in adhesin genes and genes encoding the Lap secretion system outer membrane pore component. Statistical tests: hypothesis testing was conducted in Graphpad Prism 8 using two tailed t tests with Holm-Sidak’s multiple comparison correction. P values are indicated by asterisks as follows: * P < 0.05, ** P < 0.01, *** P < 0.001, ns P > 0.05.

Next, we assessed the ability of LapA to be secreted in the absence of LapE (Figure 5B). The Δ*lapE*Δ*mapA* mutant formed a biofilm that was not different from that formed by the Δ*lapE*Δ*mapE* mutant. This observation suggests that, while MapA may utilize some portion of the LapBCE T1SS, LapA is likely not secreted through the MapBCE T1SS. Additionally, it was observed that the deletion of the *lapE* gene resulted in a greater decrease in biofilm than the deletion of the *lapA* gene. This supports a model in which MapA is capable of being secreted through both secretion systems and is therefore able to partially suppress the impact of the loss of LapA on biofilm formation. Alternatively, the differential expression of LapA and MapA may also explain the inability of LapA to use the Map secretion machinery.

### *lapA* and *mapA* gene expression is c-di-GMP responsive

c-di-GMP has been described as a common biofilm promoting factor in many bacteria (51–56). One mechanism by which this promotion of biofilm formation occurs is through the c-di-GMP-dependent transcriptional regulation of biofilm matrix components (55, 57–60). We therefore reasoned that c-di-GMP may play a role in the transcriptional regulation of adhesin genes in *P. fluorescens*.

To test the hypothesis that genes involved in the regulation of LapA and MapA production or localization are under transcriptional regulation of the c-di-GMP signaling network, we assessed the expression of these genes using the Nanostring nCounter system, which directly measures RNA transcripts produced under a given condition (61). Expression during growth on K10-T agar plates was measured in WT *P. fluorescens* Pf0-1 and two mutants with differing c-di-GMP levels: a low c-di-GMP strain lacking four diguanylate cyclases (DGCs) previously demonstrated to be important for biofilm formation, and designated here Δ4DGC (62), in which the level of c-di-GMP is roughly one third that observed in WT *P. fluorescens* Pf0-1 (63), and a high c-di-GMP mutant expressing a DGC with a mutation in its autoinhibitory site that leads to constitutive activity and a roughly 25-fold higher concentration of c-di-GMP than the concentration in WT *P. fluorescens* Pf0-1 cells, this strain is designated Δ4DGC p*gcbC*-R366E (63).

Expression of the two adhesin genes: *lapA* and *mapA*, was found to differ dramatically. Normalized *lapA* transcript counts were found to be ∼17-fold higher than *mapA* counts in WT *P. fluorescens* Pf0-1 (Figure 6A, Table 1). In addition, the *lapA* transcript was found to yield the highest transcript count compared to any of the other biofilm-related genes analyzed. Compared to the expression in WT cells, no significant changes in expression for any of the genes assessed was observed in the low c-di-GMP strain (Δ4DGC). Conversely, when expression was assessed in the high c-di-GMP variant, Δ4DGC p*gcbC* R366E, an increase in expression of all genes except for *lapD* and *lapG* was observed. The largest magnitude change in expression was observed for the *mapA* gene at ∼ 11-fold increase between the low and high c-di-GMP strains (Figure 6B, Table 1).

**Table 1.** Normalized read counts of genes reported in Figure 6A as assessed by Nanostring.

| Gene | WT | Δ4DGC | Δ4DGC *pgcbC* R366E | Fold increase in high Vs<br>WT c-di-GMP |
| --- | --- | --- | --- | --- |
| *lapA* | 88.99 (6.68) | 86.07 (2.05) | 149.07 (6.79) | 1.68 (0.08) |
| *lapB* | 12.23 (1.45) | 14.08 (0.76) | 23.01 (1.90) | 1.88 (0.16) |
| *lapC* | 9.95 (1.99) | 11.04 (0.11) | 24.54 (0.13) | 2.47 (0.01) |
| *lapE* | 9.33 (1.06) | 6.46 (0.93) | 57.41 (6.12) | 6.15 (0.66) |
| *mapA* | 5.13 (0.84) | 4.57 (0.67) | 54.50 (2.87) | 10.63 (0.56) |
| *lapD* | 11.53 (1.84) | 10.88 (0.66) | 9.83 (0.46) | 0.85 (0.04) |
| *lapG* | 5.47 (0.85) | 5.83 (0.54) | 5.06 (0.53) | 0.93 (0.10) |

<sup>a</sup>Data presented as means. Standard deviation is given in parentheses.

**Figure 6.**
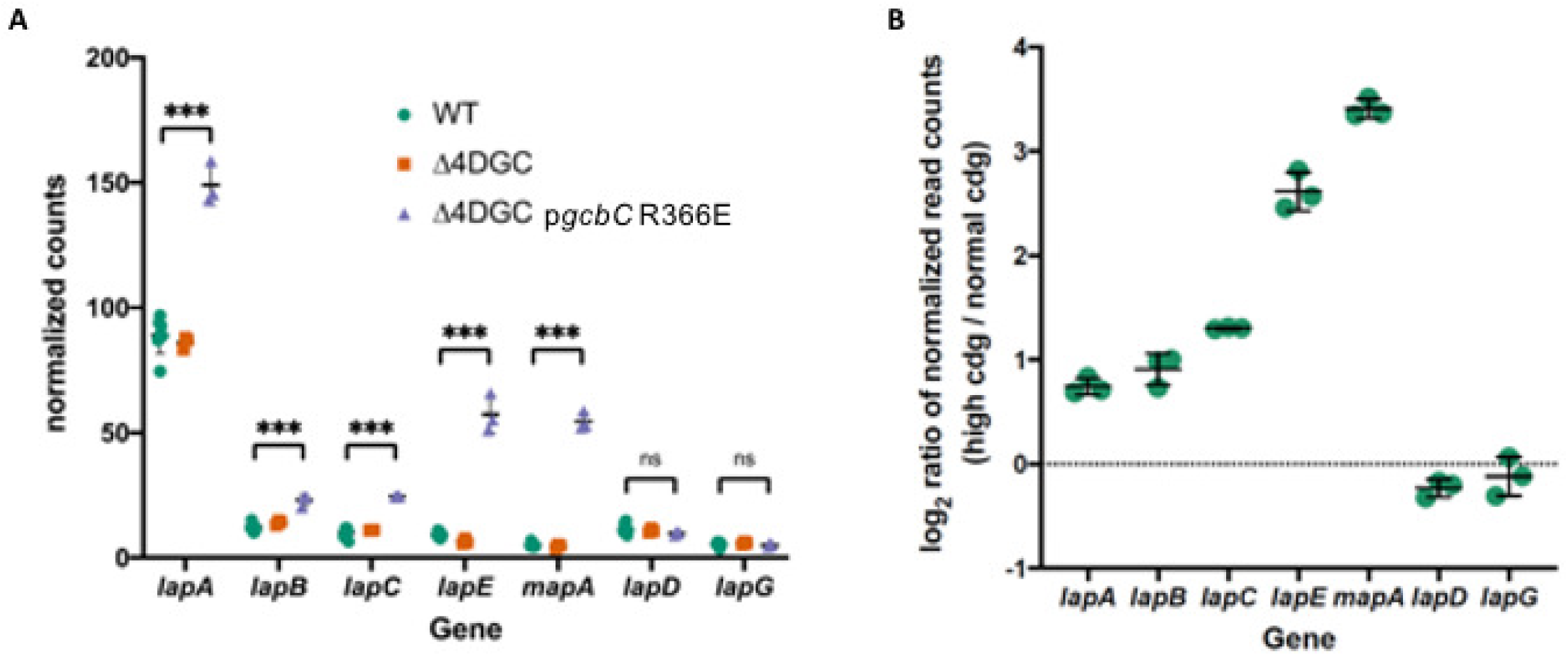
Impact of c-di-GMP levels on *lapA* and *mapA* genes expression. (A) Shown are the normalized read counts for the indicated genes in the WT, a strain with low c-di-GMP levels (D4DGC) and high c-di-GMP levels (p*gcbC* R336E). Reads were normalized to 1000 total counts across all genes, meaning that *lapA* transcript counts in *P. fluorescens* Pf0-1 cells have a mean count number of 88.99, corresponding to 8.899% of the total reads detected across all genes in that sample. Details of the method to determine read counts of expression after normalization is described in the Materials and Methods. (B) Log_2_ transformed ratios of expression for each gene indicated in the Δ4DGC p*gcbC*-R366E strain versus the WT. Statistical tests: hypothesis testing was conducted using Dunnet’s multiple comparisons test in Graphpad Prism 8 software. *** P < 0.001, ns P > 0.05.

### LapA and MapA are both required for the typical three-dimensional structure of the biofilm

We next sought to elucidate the functional significance of encoding two large adhesins in the context of the formation of a mature biofilm using a flow cell system. After 96 hours of growth in the microfluidic chamber, the WT forms a thick biofilm that fills most of the flow cell chamber (Figure 7A). The Δ*lapA* mutant is severely deficient in biofilm formation and only small clumps of cells are observed attached to the surface (Figure 7B), as reported previously (23). The Δ*mapA* mutant appears able to attach to the surface of the chamber, but the thickness of the biofilm is reduced compared to WT (Figure 7C). Finally, the Δ*lapA*Δ*mapA* mutant resulted in a biofilm comprised of sparsely distributed clumps of cells (Figure 7D). Together, these clear deficiencies in biofilm structure support a role for both adhesins in the formation of the biofilm.

**Figure 7.**
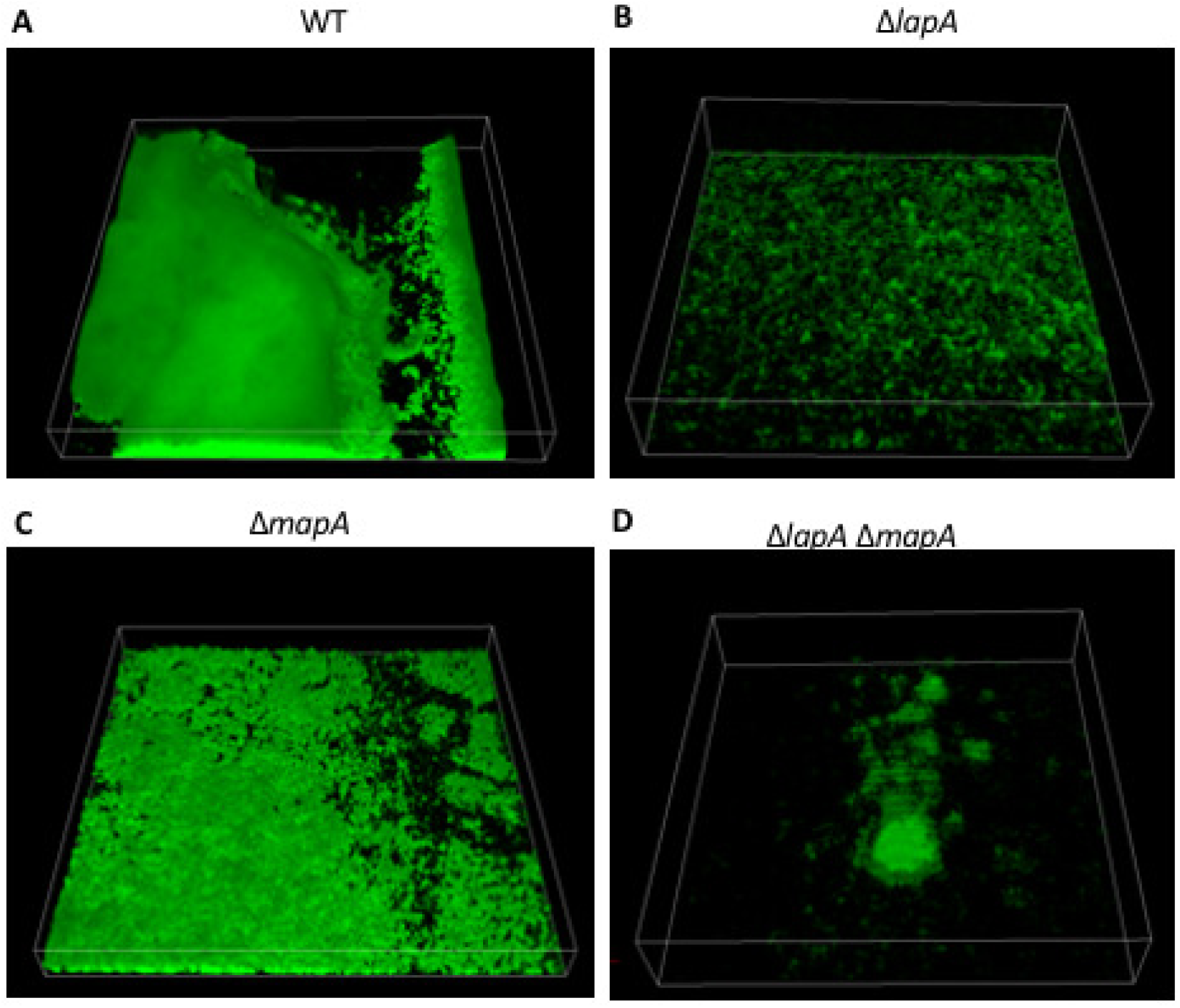
Confocal spinning disk microscopy images of biofilm formed by adhesin mutants in flow cells. The indicated strains were grown as described in the Materials and Methods and incubated at room temperature under constant flow for 96 hours before imaging. Shown is a representative image taken from one of two biological replicates.

### The *lapA* and *mapA* genes are differentially expressed in a *P. fluorescens* Pf0-1 biofilm

Given the differential impact on biofilm formation for the *lapA* and *mapA* mutants, we hypothesized that LapA and MapA may be produced in different regions of a biofilm. In order to test this hypothesis, we constructed a reporter strain in which the promoter of each adhesin is driving the expression of a different fluorescent protein at a neutral site on the chromosome, while the native loci remain unchanged and thus functional. In this strain, the promoter of the *lapA* gene drives expression of GFP while the promoter of the *mapA* gene drives production of mRuby. Thus, this strain produces LapA and MapA from their native loci, while the GFP and mRuby constructs function as reporters of the activity of the *lapA* and *mapA* promoters, respectively. Figure 8A depicts a graphical representation of this adhesin reporter construct.

**Figure 8.**
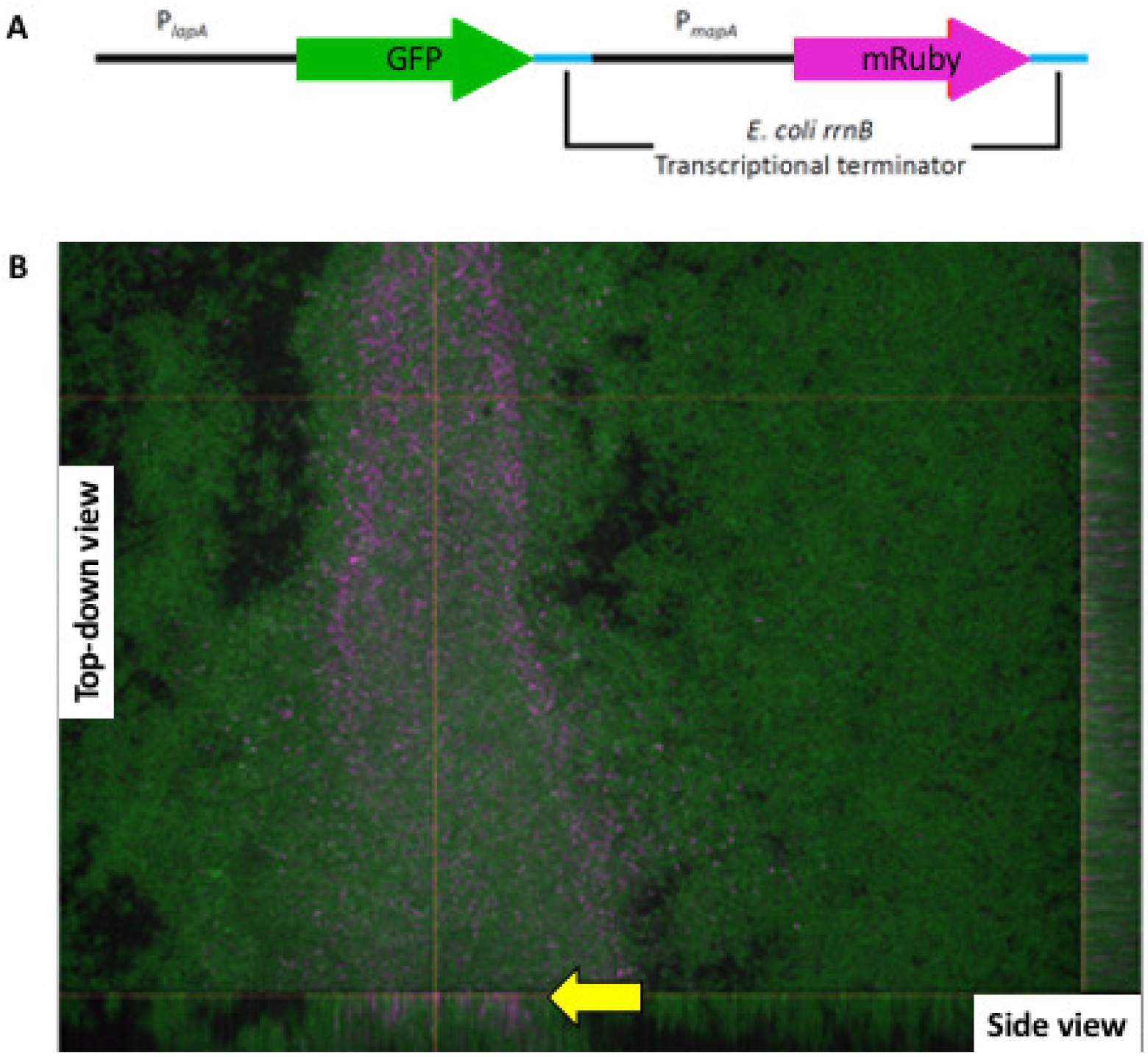
Differential expression of *lapA* and *mapA* genes in a biofilm. (A) Schematic of adhesin reporter construction indicating promoter and the gene encoding the fluorescent protein gene. P_*lapA*_ is driving the expression of the green fluorescent protein gene and P_*mapA*_ is driving the expression of the mRuby gene. Both fluorescent protein genes have a transcriptional terminator of the *rrnB* gene from *E. coli* downstream of the genes to prevent transcriptional read through. (B) Confocal image of a biofilm grown in a microfluidic device. Image taken of 96-hour biofilm with 40X oil-immersion objective lens. Green is the GFP channel excited by 488nm laser and indicate *lapA*-expressing cells. Magenta is the mRuby channel excited by 560nm laser and indicates *mapA*-expressing cells. Shown is a representative image taken from one of two biological replicates.

We grew this adhesin reporter strain in microfluidic devices under constant flow in a variant of the KA medium (see Materials and Methods for details) for 96 hours, and then imaged the biofilms formed by spinning disk confocal microscopy in order to assess expression of the *lapA* or *mapA* reporters (Figure 8B). Figure 8B shows a typical field of view in which a thick region of biofilm can be seen spanning from the bottom to the top of the center of the field of view, while the regions on either side of this central area of dense biofilm are visibly more porous and thinner. We found that only *lapA* is expressed in the areas where biofilm is thin or less dense (Figure 8B, green, right portion of the field of view), whereas robust *lapA* and *mapA* expression could both be detected in regions where the biofilm is thicker and more dense (Figure 8B, magenta). The expression of the *mapA* gene is localized to this dense biofilm region and is not observed in the more porous regions of the biofilm. In addition, *mapA* expression seemed to be localized to the region closest to the point of attachment to the glass cover slip as shown by the side projections on the right and the bottom of Figure 8B (yellow arrow).

The observations that LapA and MapA seem to be capable of supporting biofilm formation under different growth conditions and are both required for proper three-dimensional biofilm structure supports a model wherein these two adhesins play different roles during biofilm formation. However, it is not clear from these data whether those different roles are due solely to differences in transcriptional regulation and that LapA and MapA are otherwise functionally redundant, or alternatively, that the proteins have distinct functions/roles in the context of biofilm formation. To distinguish between these possibilities, we created a strain in which *mapA* expression is placed under the control of the *lapA* promoter, referred to as “Pswap”. The region described here as the *lapA* promoter is defined as spanning from the base adjacent to the *lapE* open reading frame (ORF) to the base adjacent to the *lapA* ORF (see Figure 4). The region described as the *mapA* promoter region is the equivalent region spanning from the base adjacent to the *mapE* ORF to the base adjacent to the *mapA* ORF (see Figure 4). We reasoned that this construct would lead to both *mapA* and the *mapBCE* genes encoding its cognate T1SS being transcriptionally regulated in a similar fashion to *lapA* and its cognate T1SS, and therefore allow us to make direct comparisons between the level of biofilm formed by WT and Δ*lapA* cells and that formed by the Pswap and Δ*lapA* Pswap mutants.

We tested biofilm formation of WT, Δ*lapA*, Pswap, and Δ*lapA* Pswap when grown in KA media for 6 and 16 hours (Figure 9A-B). If MapA were functionally equivalent to LapA then we would expect the Δ*lapA* Pswap mutant to behave like a complemented *lapA* mutant and form approximately WT levels of biofilm. Instead we observed no increase in biofilm formation between the Δ*lapA* and Δ*lapA* Pswap mutants at either timepoint (Figure 9A-B).

**Figure 9.**
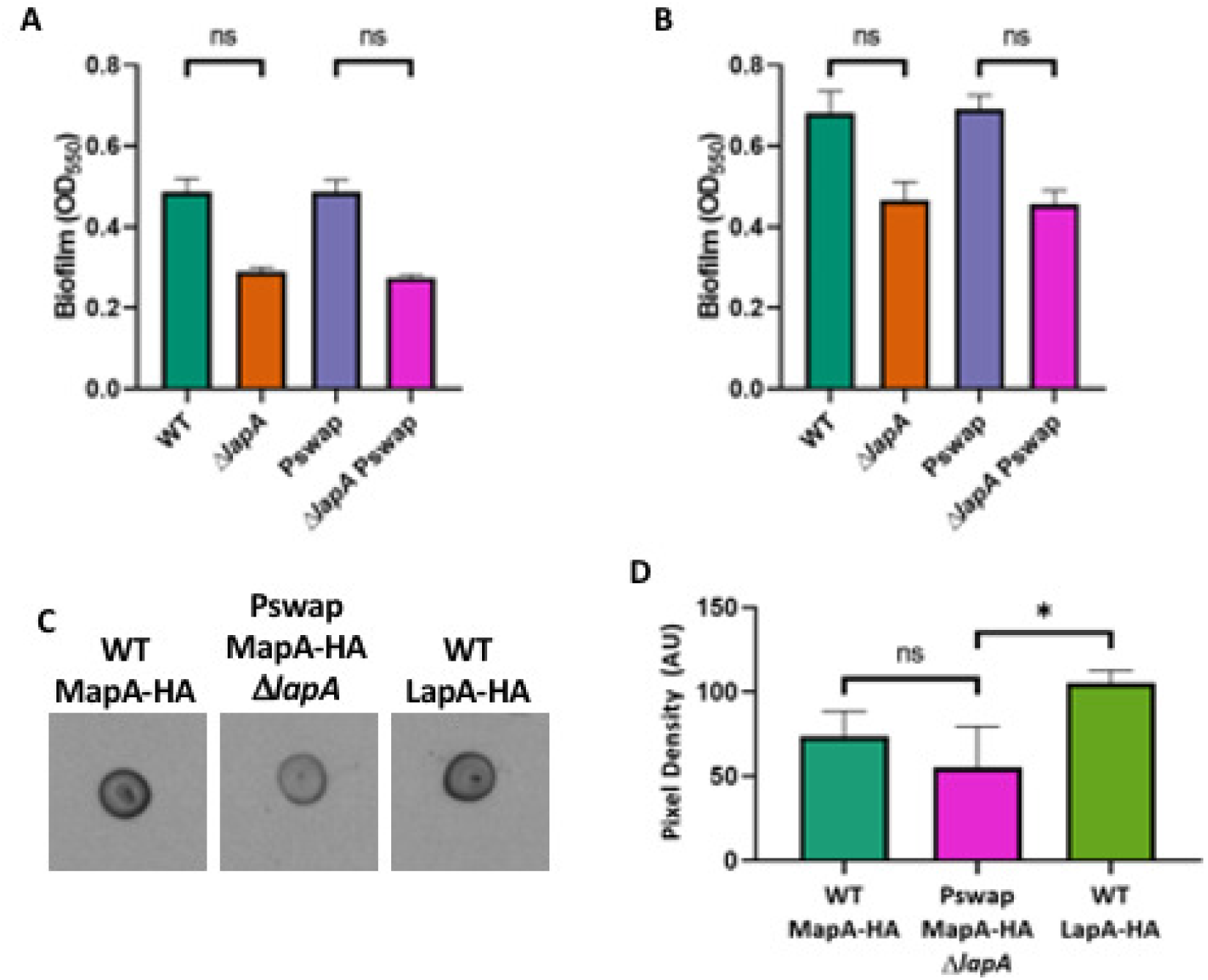
Expressing *mapA* under control of the *lapA* promoter does not rescue loss of LapA function. Quantification of biofilm formed by WT *P. fluorescens* Pf0-1, Δ*lapA*, Pswap, and Δ*lapA* Pswap after (A) 6 hours of growth in KA medium and (B) 16 hours of growth in KA medium. (C) Cell surface expression of MapA in the WT and Pswap strain. The cell surface expression of LapA is shown as an additional control. Shown is one representative experiment of three biological replicates. (D) Quantification of the cell surface expression of HA-MapA in the WT and Pswap strain, and HA-LapA in the WT strain, as indicted. Quantification was performed as decribed in the figure legend of Figure 3. Statistical tests: All hypothesis testing was conducted using two tailed t test. * P < 0.05, ns P > 0.05.

To confirm that MapA was expressed, and more importantly, surface localized in the Δ*lapA* Pswap strain, we performed a dot blot experiment as described above. The Δ*lapA* Pswap strain did indeed yield cell-surface HA-MapA, although the level was approximately 25% less than observed for the level of HA-MapA produced by the WT strain (Figure 9C-D, compare left and center of panel C), there was no significant difference between the level of MapA produced by the WT and the Pswap strain. As an additional positive control, we also blotted for the cell-surface levels of LapA in a strain carrying the HA-LapA-expressing construct (Figure 9C, right). Given that we observed no change in biofilm formation between the *lapA* and Δ*lapA* Pswap (Figure 9A-B), and the level of MapA in the Pswap strain was ∼75% that of the WT strain but not significantly different, the simplest explanation for these observations is that MapA cannot functionally substitute for LapA.

### Lap-like systems are broadly distributed throughout the Proteobacteria

The use of multiple adhesins in biofilm formation has been described in other organisms, for example BapA and SiiE in *Salmonella enterica* (64–66) and LapA and LapF in *P. putida* (45, 67–69). We were therefore interested to assess how widespread this phenomenon may be throughout the bacterial domain in the context of LapA-like adhesins. To assess the distribution of these proteins, we modified a previously described approach to identify LapD and LapG homologs in other organisms and then examined those genomes for evidence of RTX adhesins that may be targets of the LapG homolog in the same organism (29)(See Materials and Methods for a description of this approach).

Using this approach, we identified 3740 unique species encoding both LapD and LapG homologs. We next retrieved all available complete, annotated protein coding sequences for organisms of LapD and LapG encoding organisms that are available on the NCBI GenBank database. The phylogenetic class classification of these organisms is indicated in Table 2. The vast majority of these organisms whose genome encodes a LapA-like protein are members of the Proteobacteria, with most of those being members of the Gammaproteobacteria. The observation that we identified more organisms that encode putative LapD and LapG homologs in the Gammaproteobacteria than in other Proteobacteria is not necessarily indicative of an enrichment of these protein in that class as only 12% of Gammaproteobacteria represented in the GenBank database were found to encode LapD and LapG homologs. Indeed, 53% of the Zetaproteobacteria and 42% of the Hydrogenophilalia were found to encode both LapD and LapG, indicating that this system may be more highly represented in classes other than the Gammaproteobacteria. However, we identified almost no LapD and LapG homologs outside of the Proteobacteria (Table 2), indicating that these proteins may be largely restricted to this phylum.

**Table 2.**
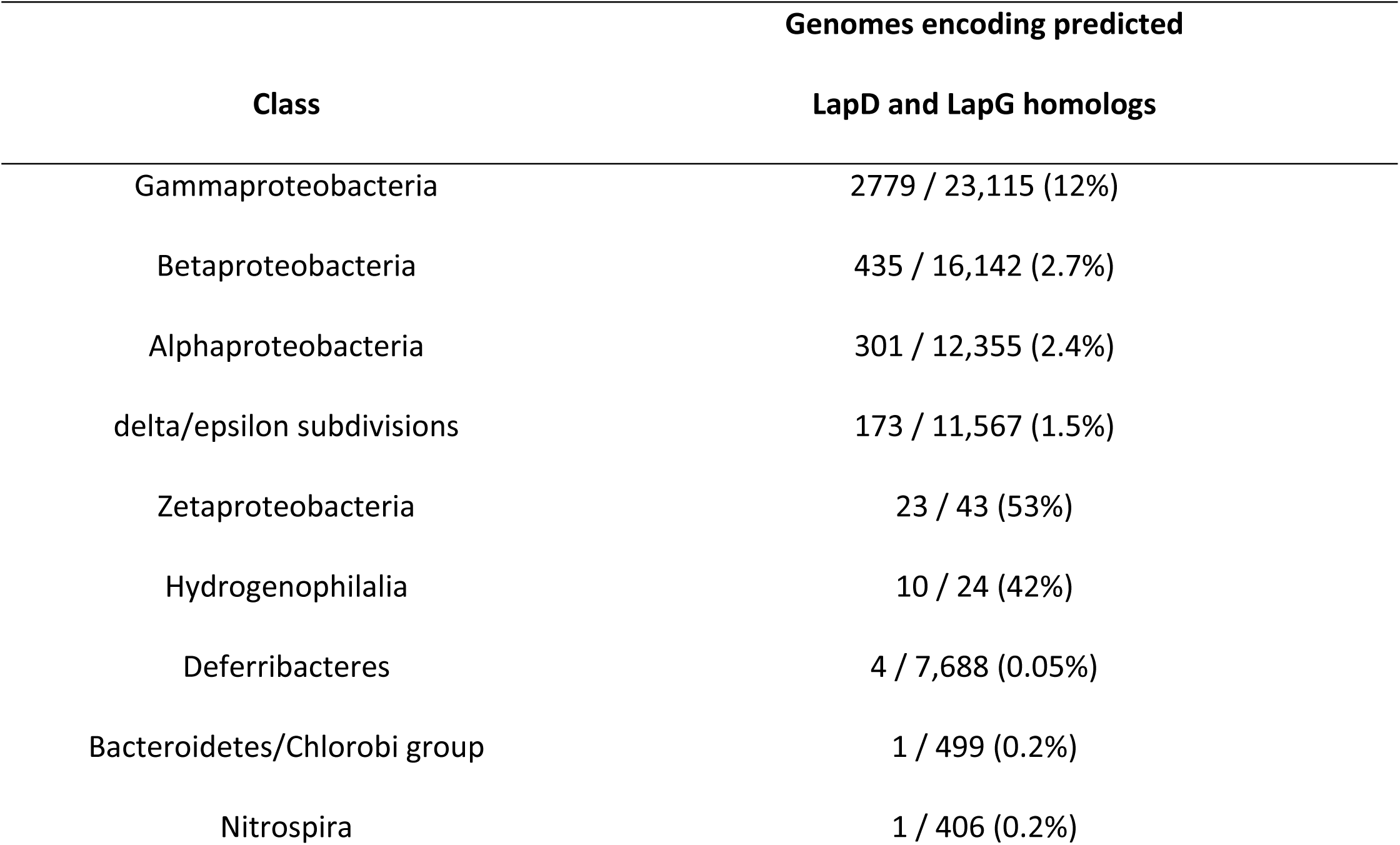
Number of genomes found to encode both putative LapD and LapG homologs in each of the given taxa.

Having collected a list of LapD and LapG encoding organisms, we next retrieved amino acid sequences of all annotated genes in these organisms and searched these sequences for features that identify them as putative LapA-like proteins (see Materials and Methods for details). Briefly, the criteria for this characterization are: (i) having a size in excess of 1000 amino acids; (ii) the presence of putative RTX repeats in the C-terminal portion of the protein; and, (iii) the presence of a putative LapG cleavage site in the N-terminal portion of the protein. Through this search we identified 1020 organisms encoding predicted LapA-like proteins in addition to LapG and LapD homologs. Among those organisms, approximately one quarter were found to encode multiple LapA-like adhesins (Table 3). Interestingly, while many LapD- and LapG-encoding organisms encoded proteins that could be identified as LapA-like proteins, 2720 organisms in our dataset were found to encode LapD and LapG homologs, but were not found to encode proteins that we could readily identify as LapA-like using our criteria, which could be due to the lack of such proteins or the difficulty in annotation of such large adhesins.

**Table 3.** Number of putative LapA-like adhesins predicted to be encoded in genomes in which putative LapD and LapG homologs were identified.

| Number of adhesins |  | Number of organisms |
| --- | --- | --- |
| 1 |  | 352 |
| 2 |  | 109 |
| 3 |  | 20 |
| 4 |  | 3 |

### Pseudomonads encodes LapA-like proteins with a distinct size distribution

RTX-adhesins are typically large and are often the largest proteins produced by the bacteria in which they are found. Most of the length of these proteins is composed of tandem repeats. These repeats have in some cases been described to function as extender domains which hold the adhesive C-terminal domains far away from the cell surface (70, 71). Thus, while the functions of the various domains of LapA-like proteins are still in most cases unclear, the protein length may be an important characteristic. We therefore sought to assess the distribution of LapA-like protein sizes found within the three most highly represented genera in our dataset: *Vibrio, Shewanella*, and *Pseudomonas* (Figure 10). By plotting the distribution of LapA-like proteins we can assess the sizes of adhesins encoded by organisms containing one or multiple such proteins.

**Figure 10.**
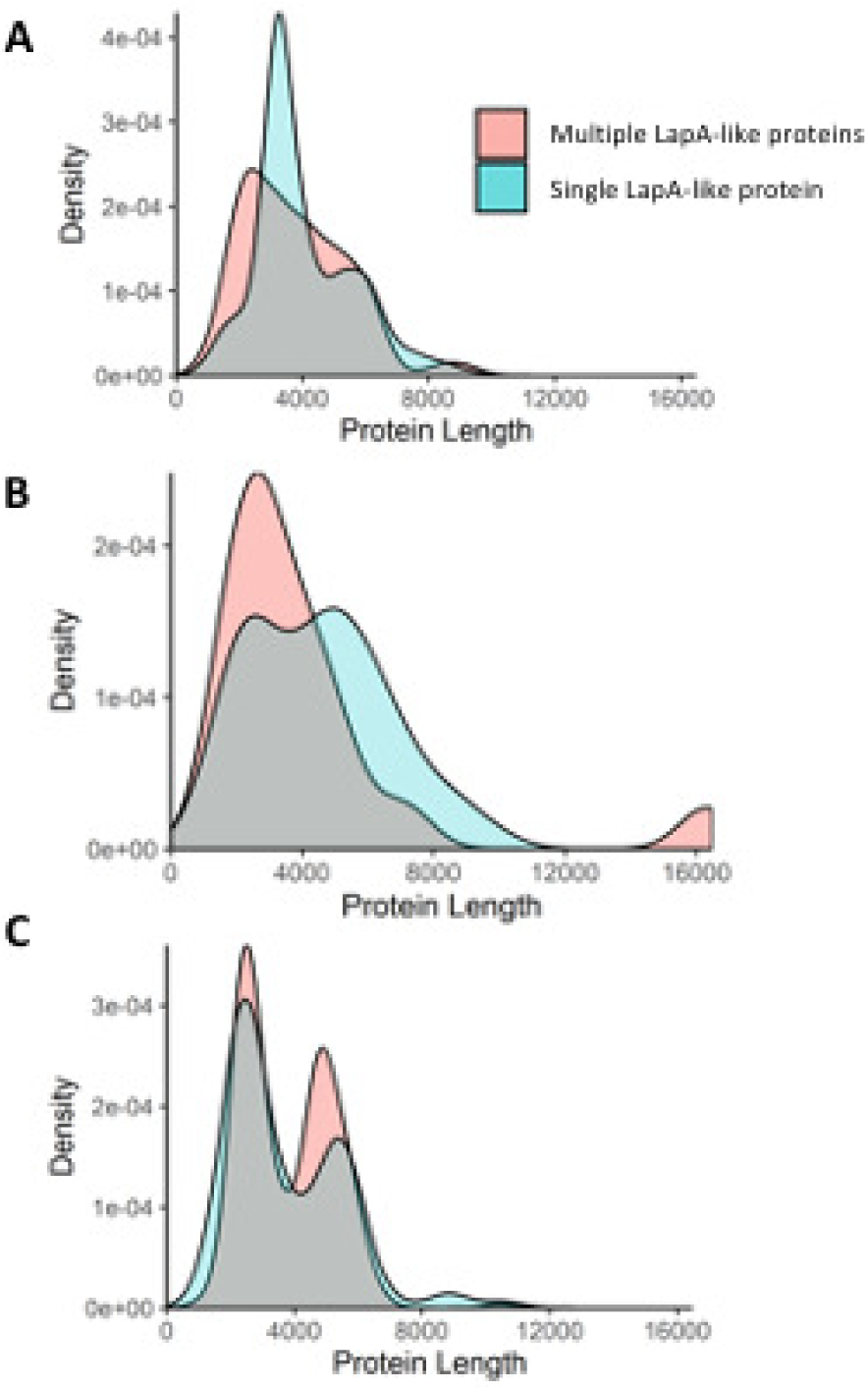
Size distribution of LapA-like proteins encoded by selected genera. All organisms found to encode both LapD and LapG, and putative LapA-like proteins were grouped based on whether they are predicted to encode a single, or multiple LapA-like proteins. We then assessed the distribution of the protein sizes of adhesins encoded by organisms with a single (blue shading) or multiple adhesins (orange shading). Shown are density plots representing the size distributions of LapA-like proteins found in the following genera: (**A**) *Vibrio*, (**B**) *Shewanella*, and (**C**) *Pseudomonas*.

LapA-like proteins encoded by *Vibrio* species with just one such protein exhibit a striking peak at a length of around 3000 amino acids and a shoulder at around 5000 amino acids (Figure 10A). The major peak of organisms encoding a single LapA-like protein corresponds to a large number of *V. cholerae* isolates, including *V. cholerae* O1 biovar El Tor that encode an almost identical adhesin, while the shoulder at around 5000 amino acids corresponds to a variety of different species. Interestingly, there is some diversity in the number and size of LapA-like proteins encoded by *V. cholerae* isolates, with some encoding two and others just one such protein. In addition, there appear to be several distinct LapA-like variants encoded among isolates of *V. cholerae*. Some of these variant proteins appear to share very little sequence similarity to one another, while others appear to be mostly identical to another variant, differing mostly due to a large indel. For example, the LapA-like protein encoded by the *V. cholerae* O1 biovar El Tor is highly similar to a LapA-like protein encoded by *V. cholerae* TMA 21, but the El Tor protein is ∼880 amino acids shorter. How these putative LapA-like proteins differ functionally from one another is unclear. However, the variation seen within the *V. cholerae* species may indicate that this protein is important for adaptation to different lifestyles of these organisms.

Within the genus *Shewanella*, no one species is represented dramatically more than any other in our dataset. Accordingly, we do not see a dramatic peak in the protein size distribution corresponding to the adhesin encoded by a single species. Most of the species within this genus encode proteins that differ from those of other species both in size and at the sequence level, leading to a smooth curve when the size distribution is plotted (Figure 10B). Interestingly, the largest LapA-like protein in our dataset is encoded by *Shewanella woodyi* ATCC 51908 by the gene Swoo_0477. This protein is predicted to be 16,322 amino acids, which is over 5000 amino acids longer than the next longest protein we identified as a LapA-like protein. While we are not aware of any experimental evidence to support the enormous size of this protein, it is nonetheless to our knowledge the largest predicted LapA-like protein.

The distribution of protein sizes observed within the *Pseudomonas* genus has a bimodal distribution with peaks around 3000 and 5000 amino acids – the approximate sizes of MapA and LapA respectively – among both organisms that encode a single and multiple LapA-like proteins (Figure 10C). There are many isolates in our dataset that have not been assigned a species designation and which encode proteins with sizes approximately equal to that of LapA and MapA. These isolates may represent over-sampling of a small number of species that could exaggerate the observed distribution. However, there are also a considerable number of *Pseudomonas* with species designations that exhibit this pattern.

### *Pseudomonas* species encoding multiple LapA-like proteins are mostly *P. fluorescens* lineage members

Having identified which *Pseudomonas* species encode LapD, LapG and LapA-like proteins, we next sought to assess the phylogenetic distribution of these proteins within the genus. We therefore constructed a phylogenetic tree of the *Pseudomonas* species that are represented in the GenBank database (Figure 11). The phylogenetic tree presented here has a similar topology to that of trees generated through the same approach in previous studies (72, 73) (See Figure 11 legend and Methods and Materials for details).

**Figure 11.**
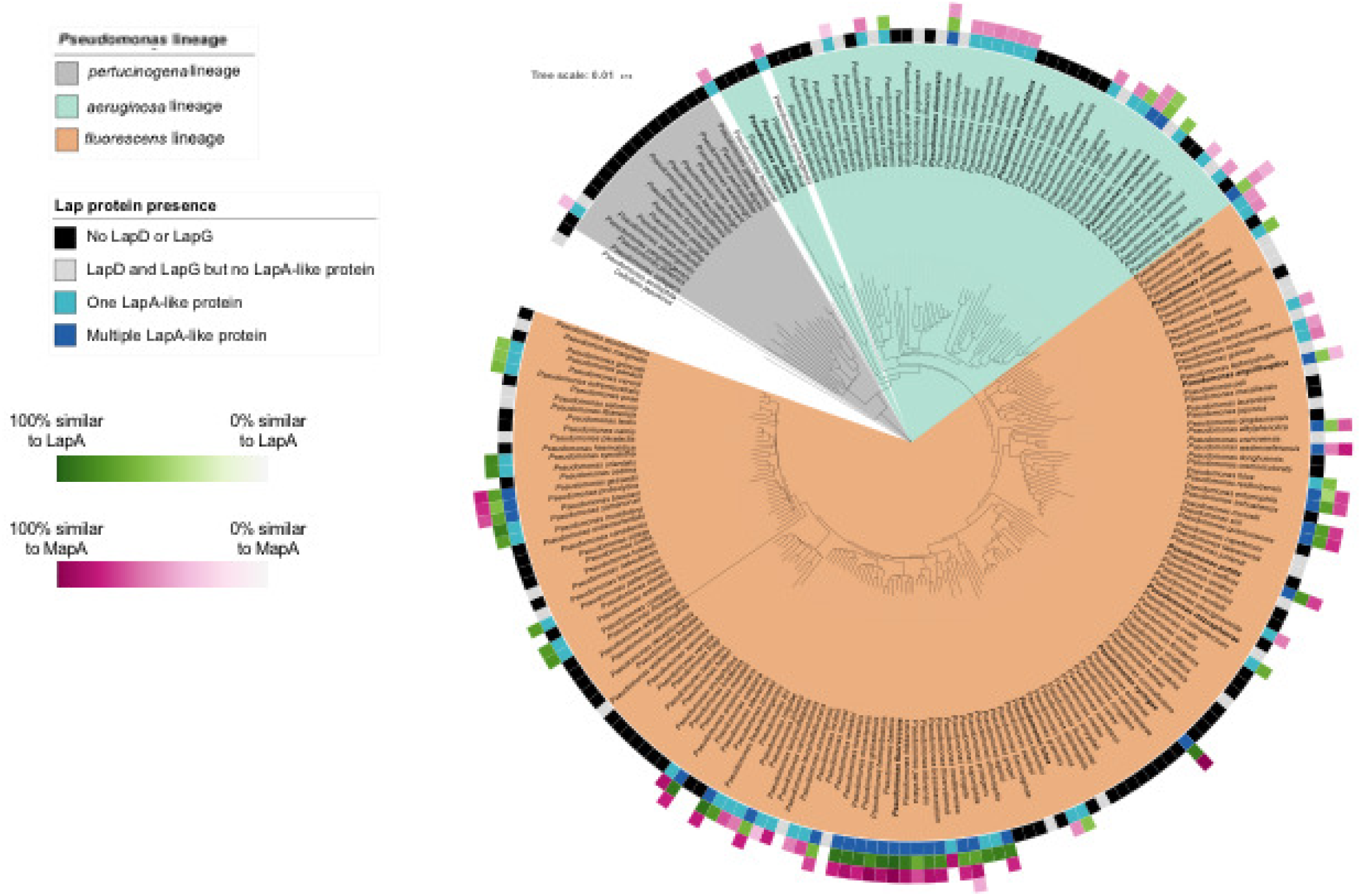
Phylogenetic distribution of the Lap proteins among members of the *Pseudomonas* genus. Shown is phylogenetic tree of *Pseudomonas* lineages. The inner-most colored ring around the tree indicates whether LapD and LapG are encoded by the indicated species (as determined by finding the intersection of the species lists retrieved from the NCBI conserved domain database for pfam06035 and pfam16448 domains) and how many LapA-like proteins were identified in that species using the analysis described in the Materials and Methods. The outer-most 3 rings indicate the percent similarity between the LapA-like proteins identified in each species and either LapA or MapA. Amino acid sequence of putative LapA-like proteins were aligned with LapA and MapA in pairwise alignments using MUSCLE (74) (https://github.com/GeiselBiofilm/Collins-MapA/tree/master/Supplemental_File_Alignments) and the percent sequence similarity was determined by counting the positions identified as similar by the MUSCLE program. The percent similarity of whichever was most similar (either LapA or MapA) is indicated with darker color indicating higher similarity. The order of the three rings indicating similarity with LapA and MapA indicates the relative size of the LapA-like proteins encoded by each organism. The inner-most of the three rings is the largest LapA-like protein encoded by that organism. The second and third rings represent alignments of the second- and third-largest LapA-like proteins identified in those organisms that are predicted to encode multiple LapA-like proteins. The MLSA phylogenetic tree here shows the clustering of representatives *Pseudomonas* species found in the GenBank database based on the analysis of concatenated alignments of 16S rRNA, *gyrB, rpoB*, and *rpoD* genes. Distance was calculated using the Jukes-Cantor model and the tree was constructed using neighbour-joining. The *P. aeruginosa* (light blue-green shading), *P. fluorescens* (orange shading), and *P. pertucinogena* (grey shading) lineages described by Mulet et al. (73) are highlighted with the indicated colors. The namesake species of the groups described by García-Valdés and Lalucat (95) are in bold font. The *P. aeruginosa* and *P. fluorescens* lineages described by Mulet et al. (73), and the *P. pertucinogena* lineage described by Peix et al. (72) are also apparent in our tree and are indicated using light blue-green, orange, and grey shading, respectively in Figure 11 (colored according to the lineages described by Peix et al. (72)). The most notable difference between the tree presented here and previously reported trees is the presence of a distinct *P. luteola* and *P. duriflava* clade, while Peix et al. (72) presented them as part of the monophyletic *P. aeruginosa* lineage. However, the placement of *P. luteola* within the *aeruginosa* lineage is not strongly supported. The study by Mulet et al. (73) placed *P. luteola* outside of the *P. aeruginosa* lineage and the bootstrap support for Peix et al. (72) positioning the *P. luteola* and *P. duriflava* clade within the *P. aeruginosa* lineage is less than 50%. Three other species were not present in previously published phylogenies of *Pseudomonas* and do not fit neatly into the three lineages described previously: *P. pohangensis, P. hussainii*, and *P. acidophila*. Both *P. pohangensis* and *P. hussainii* are positioned between the *P. pertucinogena* lineage and the other two lineages. *P. acidophila* is positioned outside of the rest of the genus, consistent with a recent study that argued that *P. acidophila* should be reclassified as *Paraburkholderia acidophila* (96). While the *P. pertucinogena* lineage described by Peix et al. (72) is well supported by the tree we present here, the species *P. pertucinogena* is absent from our tree. This is because we only included species present in the GenBank database in our tree. At time of writing there is an available genome sequence record of *P. pertucinogena* (Bioproject accession PRJNA235123), but the absence of an assembled genome from GenBank means that the organism was not included in our analysis of the distribution of LapA-like proteins. Nevertheless, the overall tree topology is similar overall to the previously reported trees outlined above.

Through our analysis we identified LapD and LapG homologs throughout the *P. aeruginosa* and *P. fluorescens* lineages of the *Pseudomonas* genus (Figure 11, innermost ring). However, LapD and LapG homologs were only identified in 2 out of the 17 (11.7%) representatives of the *P. pertucinogena* lineage and only one of those was found to encode a LapA-like protein. Within the *P. aeruginosa* and *P. fluorescens* lineages roughly the same proportion of species were found to encode LapD and LapG (56% and 59% respectively). In addition, the proportion found to encode both LapD and LapG, but no detectable LapA-like protein was also similar between the two lineages with 9 out of 50 species (18%) in the *P. aeruginosa* lineage and 26 out of 147 species (17.7%) in the *P. fluorescens* lineage. Interestingly, these two lineages differ dramatically in the number of adhesins detected in species that were found to encode LapD, LapG, and a LapA-like protein. Within the *P. aeruginosa* lineage, the number of species encoding a single adhesin is 15 (30%) and the number encoding more than one is 4 (8%). Within the *P. fluorescens* lineage, the number encoding a single adhesin is 32 (21%) and the number encoding multiple adhesins is 29 (20%). It is therefore clear that species encoding multiple LapA-like proteins are mostly found within the *P. fluorescens* lineage.

While species encoding LapD, LapG, and LapA-like proteins are found across both the *P. aeruginosa* and the *P. fluorescens* lineages, the distribution of these proteins within those lineages is not uniform. There are multiple clades within each lineage that seem to lack identifiable homologs for any of the three proteins. For example, within the *P. aeruginosa* lineage, the clade containing *P. luteola* and *P. duriflava*, and the clade containing *P. oryzihabitans* are predominated by species that do not encode identifiable Lap proteins. Within the *fluorescens* lineage, the clade containing *P. syringae* is also predominated by species lacking identifiable Lap proteins. Interestingly, while species in the clade containing *P. syringae* mostly lack Lap proteins, *P. syringae* encodes both LapD and LapG as well as two identifiable LapA-like proteins.

### LapA and MapA share high levels of sequence similarity with LapA-like proteins encoded by other *fluorescens* lineage members

Many members of the *P. fluorescens* lineage encode multiple LapA-like proteins and the distribution of sizes of these proteins is similar to the sizes of LapA and MapA. Therefore, we hypothesized that the LapA-like proteins encoded by *Pseudomonas* species in our dataset are similar to LapA and MapA, while those encoded by organisms from other genera are less similar to LapA and MapA. To assess this idea, we conducted pairwise alignments of the amino acid sequences of every LapA-like protein that we identified in our analysis with both LapA and MapA using MUSCLE v3.8 (74) (Supplemental file 1 and https://github.com/GeiselBiofilm/Collins-MapA/tree/master/Supplemental_File_Alignments).

We found that within the *Pseudomonas* genus many organisms encode adhesins that are highly similar to LapA and MapA (Figure 11, rings of green and magenta squares). For example, *P. syringae* encodes two LapA-like proteins; the larger of the two is 87% identical and 90% similar to LapA, while the smaller of the two is 97% identical and 99% similar to MapA (Figure 11, supplemental file 1). Interestingly, most of the organisms within the *P. fluorescens* lineage that encode multiple adhesins were found to encode proteins with very high similarity with LapA and MapA. However, LapA-like proteins encoded by members of the *P. aeruginosa* lineage have relatively lower similarity to LapA and MapA proteins more broadly.

Outside of the *Pseudomonas* genus the occurrence of proteins with very high similarity to LapA or MapA is lower. However, many organisms do encode proteins with moderate similarity (over 50% across the entire length of the protein) with LapA or MapA. In some cases, the similarity between these proteins is mostly localized to the N- and C-terminal portions of the proteins which contain the anchor domain and RTX repeats, respectively, which are characteristic of LapA-like proteins (75, 76). In other cases, it appears that similarity between adhesins is localized to the central portion of the protein. For example, the LapA-like protein encoded by *Aeromonas salmonicida* has 38% amino acid identity (68% similarity) to MapA across the whole length of the protein. However, there is a highly similar region of these proteins located in the first half of their central portion (54% identity, 85% similarity between positions 393 – 1620), while the N- and C-terminal portions of the proteins have relatively low sequence identity (https://github.com/GeiselBiofilm/Collins-MapA/tree/master/Supplemental_File_Alignments). This pattern is contrary to the typically high conservation of the N- and C-termini and low conservation of the central portion of LapA-like proteins (75, 76).

## Discussion

In this manuscript we present MapA, a second type 1-secreted, large RTX adhesin that contributes to biofilm formation by *P. fluorescens*. We demonstrate that MapA is surface localized and can contribute to biofilm formation in the absence of LapA, at least under one growth condition we have identified to date. We also present genetic data showing that MapA surface localization is likely controlled by the previously described c-di-GMP effector protein, LapD and the periplasmic protease, LapG (see model in Figure 12A).

**Figure 12.**
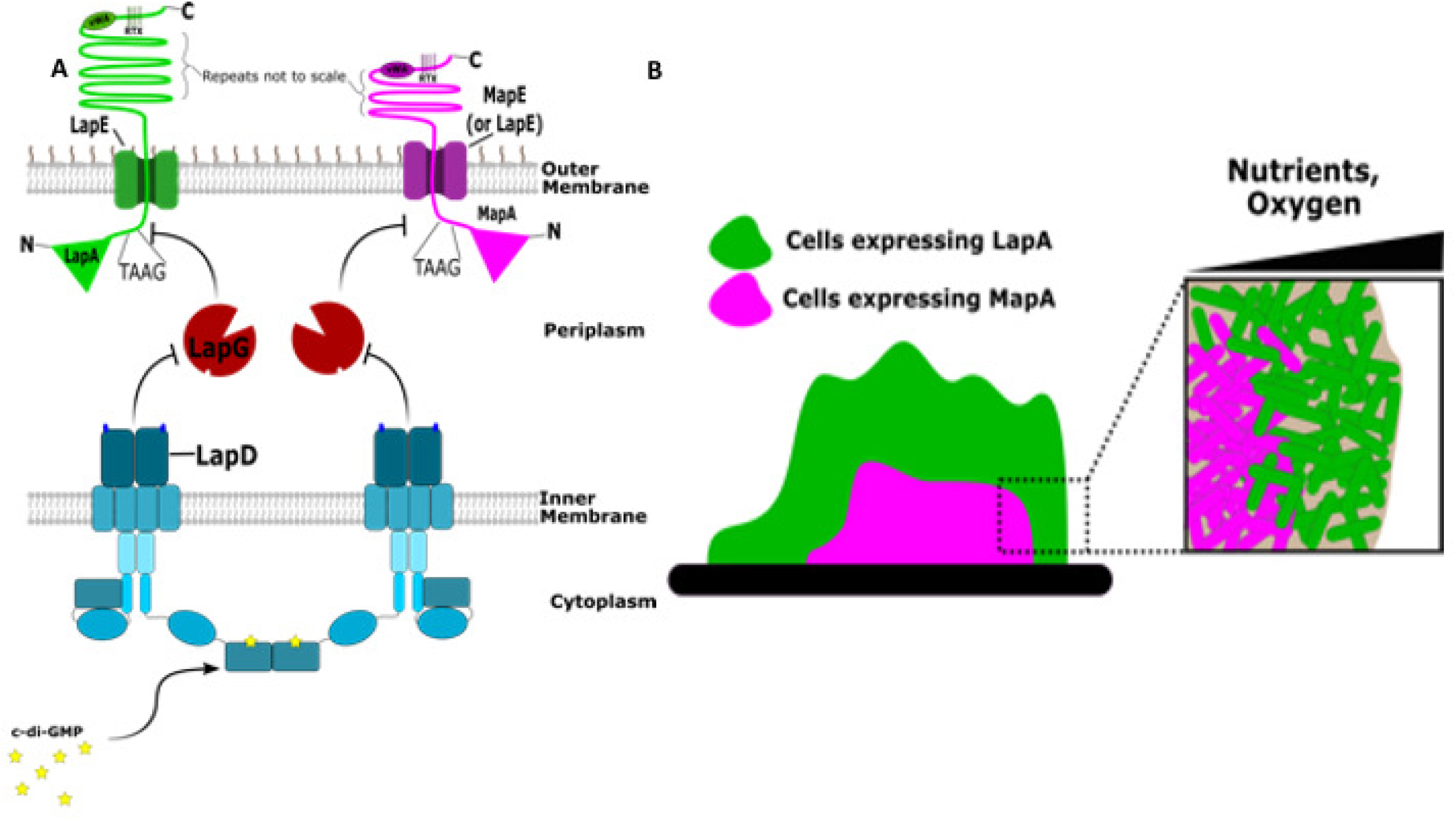
Model of the mechanisms by which LapA and MapA contribute to biofilm formation. (A) Shown is the model for c-di-GMP regulation of cell surface localization of LapA and MapA. Representations of the LapA (left) and MapA (right) adhesins anchored in the outer membrane via LapE and MapE, respectively. Also shown is the periplasmic protease LapG and the inner membrane, c-di-GMP receptor LapD which regulates LapG activity. (B) Representation of the expression of *lapA* and *mapA* in a biofilm. Indicated are possible roles for nutrient and/or oxygen limitation in controlling the expression of the *mapA* gene.

Our bioinformatic analysis indicates the distribution of similar multiple-adhesin systems throughout the Proteobacteria. Through our analysis we identified many LapD and LapG homolog-encoding organisms that encode differing numbers of putative LapA-like proteins. However, we also identified many organisms that seem to encode LapD and LapG homologs, but no identifiable LapA. Even within the *Pseudomonas* genus, we identified multiple species that seemed to lack any identifiable LapA (Figure 11). It is possible that LapA-like proteins are present in these organisms and that our approach was not able to identify them or that the protein sequence database that we used as the source of our data is incomplete due to misannotation of these large, often repeat-rich proteins. In addition, the search parameters we used to identify LapA-like proteins are based on the characterization of small number of proteins characterized to date (76), so it is possible that we have designed our search in such a way that some authentic LapA-like proteins were excluded. However, in the case of *P. aeruginosa*, a LapD and LapG homolog are present but no LapA-like protein exists. Instead, LapG is able to proteolytically process a non-canonical target CdrA and regulate its localization to the cell surface (77). CdrA is secreted through a type Vb secretion process and is seemingly evolutionarily unrelated to LapA. Therefore, we hypothesize that some of the organisms we have identified that lack LapA-like proteins may encode proteins that have convergently evolved to be regulated by LapG. Thus, our analysis may provide a good basis on which to identify organisms that may encode other novel LapG-regulated proteins.

Interestingly, while both the *P. aeruginosa* and *P. fluorescens* lineages contain many organisms encoding LapD, LapG, and LapA-like proteins, the *P. pertucinogena* lineage contains only two such species. The presence of LapD and LapG homologs in organisms outside of the *Pseudomonas* genus suggest that these proteins were lost in the *P. pertucinogena* lineage rather than gained in the other *Pseudomonas* lineages. *P. pertucinogena* lineage members are thought to be more niche specialized than other *Pseudomonas* species and have reduced genome size and metabolic versatility (78–81). Thus, it is possible that the genome reduction that is thought to have occurred in the *P. pertucinogena* lineage is responsible for the apparent absence of LapD, LapG and LapA-like proteins in these organisms.

Through our analysis of the sequence similarity between LapA-like proteins, we have found that many organisms both within the *Pseudomonas* genus and in other genera encode LapA-like proteins with high sequence similarity to LapA or MapA. Within the *Pseudomonas* genus these LapA-like proteins often have high sequence similarity to LapA and MapA across the whole length of the protein, particularly within the *P. fluorescens* lineage. Organisms from other genera were found to encode LapA-like proteins with lower sequence similarity with that of LapA and MapA over the whole length of the protein. However, in some cases portions of the central region of these proteins appear to be highly identical. The precise function of the diverse repeated sequences found in LapA-like proteins is poorly understood (76). In some cases an extender-like function has been described which positions functional or substrate-binding domains of the protein away from the cell surface (70, 71). However, in the case of LapA, the repeats are thought to directly bind to hydrophobic surfaces (26). It is therefore possible that LapA-like proteins with high identity over parts of their central portion may contain conserved substrate-binding domains or bind to surfaces with similar physical characteristics.

How MapA is contributing to biofilm formation is still not entirely clear. Loss of MapA function results in a defect in biofilm formation in static 96 well dish assays, at least in a select medium condition. Similarly, MapA was required for robust biofilm formation in a microfluidic device. The phenotype of the *mapA* mutants is also clearly distinct from that of strains lacking *lapA*. What is the basis for this difference in phenotype? One obvious explanation is that the LapA protein is ∼200 kDa larger than MapA, and there are clear differences in the sequence of the central repeat regions. Differential expression of the proteins might also impact their differential contributions to biofilm formation. For example, we observed differential spatial expression patterns of the *mapA* and *lapA* genes in biofilms grown in microfluidics chambers under flow. However, our studies using the Pswap construct, which allows *mapA* to be expressed via the *lapA* promoter, argues against this hypothesis. We note that *mapA* expression is only seen in large, thicker regions of biofilms, which presumably would be oxygen or nutrient limited. Thus, we postulate that oxygen, nutrients or energy status of the cells could contribute to the regulation of *mapA* gene expression (Figure 12B). Furthermore, the transcriptional data presented in this study demonstrates that increased levels of c-di-GMP led to dramatic increases in expression of both adhesin-encoding genes, with *mapA* gene expression increasing ∼11-fold in a strain with high versus low levels of c-di-GMP. This observation is consistent with the fact that loss of MapA shows a phenotype in static assays only when arginine is present, given that arginine has been show previously to stimulate c-di-GMP level in Pseudomonads (39, 41, 82). Complicating our analysis is the fact that our genetic studies indicate that MapA may utilize the Lap secretion system for surface localization. Nevertheless, in sum, our data suggest that nutrient/oxygen limitation and high c-di-GMP levels contribute positively to the elaboration of the MapA adhesin and thus its ability to contribute to biofilm formation. Studies are ongoing to further dissect the regulation of the *mapA* gene and its gene product. The observation that some Pseudomonads can express two discrete LapA-like proteins adds to the complex strategies that bacteria can utilize to establish their extracellular matrix during biofilm formation.

## Materials and Methods

### Strains and media

The strains used in this study can be found in Table 4. *P. fluorescens* Pf0-1 and *E. coli* S17 (83, 84) were used throughout this study. *P. fluorescens* and *E. coli* were grown at 30°C when containing plasmids; *E. coli* strains were otherwise grown at 37°C. Media used in this study were lysogeny broth (LB), K10T-1 (83), and two new modifications of previously used media. The first, KA, is a modification of K10-T in which the changes are the removal of glycerol and tryptone, and the addition of 0.4% w/v L-arginine HCl. The second medium was used for growing *P. fluorescens* Pf0-1 in microfluidics devices and is a modification of BMM medium previously described (85). This BMM medium was modified by reducing the concentration of glycerol and (NH_4_)_2_SO_4_ by 16-fold and adding L-arginine. The final concentration of each in this medium was therefore 0.009375% v/v glycerol, 472.5µM (NH_4_)_2_SO_4_, and 0.025% L-arginine mono-HCl (Sigma). When *E. coli* was grown harboring the allelic exchange plasmid pMQ30 or the expression plasmid pMQ72, the growth medium was supplemented with 10µg/µl gentamycin. When *E. coli* was grown harboring the plasmid pMQ56 or the helper plasmid pUX-BF13 the growth medium was supplemented with 50µg/µl carbenicillin. When *P. fluorescens* was grown harboring pMQ72, the growth medium was supplemented with 30µg/µl gentamycin. Allelic exchange was carried out as previously reported (86).

**Table 4.**
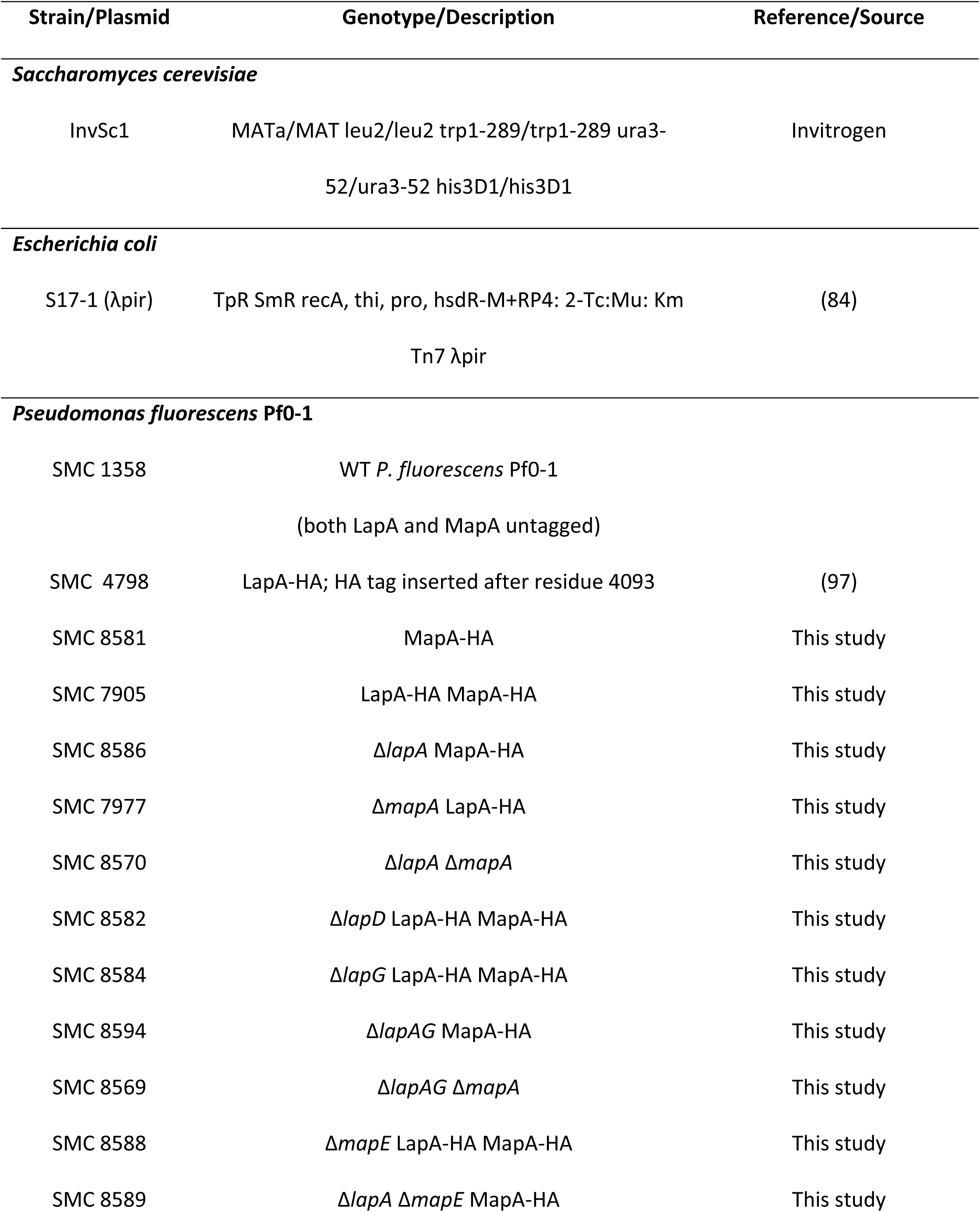
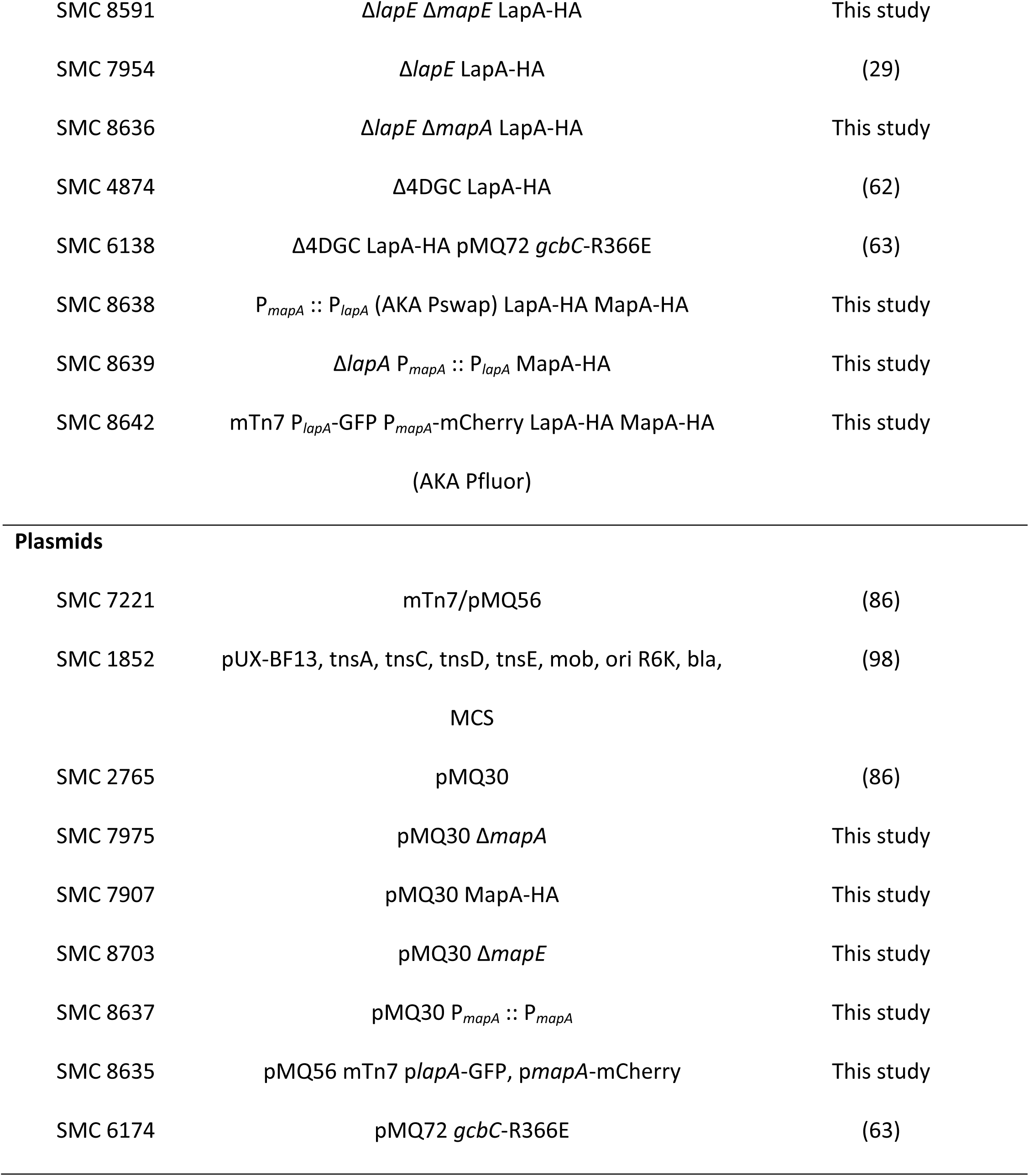
Strains and plasmids used in this study.

| Strain/Plasmid | Genotype/Description | Reference/Source |
| --- | --- | --- |
| ***Saccharomyces cerevisiae*** |  |  |
| InvSc1 | MATa/MAT leu2/leu2 trp1-289/trp1-289 ura3-52/ura3-52 his3D1/his3D1 | Invitrogen |
| ***Escherichia coli*** |  |  |
| S17-1 (λpir) | TpR SmR recA, thi, pro, hsdR-M+RP4: 2-Tc:Mu: Km<br>Tn7 λpir | (84) |
| ***Pseudomonas fluorescens* Pf0-1** |  |  |
| SMC 1358 | WT *P. fluorescens* Pf0-1<br>(both LapA and MapA untagged) |  |
| SMC 4798 | LapA-HA; HA tag inserted after residue 4093 | (97) |
| SMC 8581 | MapA-HA | This study |
| SMC 7905 | LapA-HA MapA-HA | This study |
| SMC 8586 | $\Delta lapA$ MapA-HA | This study |
| SMC 7977 | $\Delta mapA$ LapA-HA | This study |
| SMC 8570 | $\Delta lapA \Delta mapA$ | This study |
| SMC 8582 | $\Delta lapD$ LapA-HA MapA-HA | This study |
| SMC 8584 | $\Delta lapG$ LapA-HA MapA-HA | This study |
| SMC 8594 | $\Delta lapAG$ MapA-HA | This study |
| SMC 8569 | $\Delta lapAG \Delta mapA$ | This study |
| SMC 8588 | $\Delta mapE$ LapA-HA MapA-HA | This study |
| SMC 8589 | $\Delta lapA \Delta mapE$ MapA-HA | This study |
| SMC 8591 | $\Delta lapE \Delta mapE$ LapA-HA | This study |
| SMC 7954 | $\Delta lapE$ LapA-HA | (29) |
| SMC 8636 | $\Delta lapE \Delta mapA$ LapA-HA | This study |
| SMC 4874 | $\Delta 4DGC$ LapA-HA | (62) |
| SMC 6138 | $\Delta 4DGC$ LapA-HA pMQ72 *gcbC*-R366E | (63) |
| SMC 8638 | $P_{mapA} :: P_{lapA}$ (AKA Pswap) LapA-HA MapA-HA | This study |
| SMC 8639 | $\Delta lapA P_{mapA} :: P_{lapA}$ MapA-HA | This study |
| SMC 8642 | mTn7 $P_{lapA}$-GFP $P_{mapA}$-mCherry LapA-HA MapA-HA<br>(AKA Pfluor) | This study |
| Plasmids |  |  |
| SMC 7221 | mTn7/pMQ56 | (86) |
| SMC 1852 | pUX-BF13, *tnsA*, *tnsC*, *tnsD*, *tnsE*, *mob*, *ori* R6K, *bla*,<br>MCS | (98) |
| SMC 2765 | pMQ30 | (86) |
| SMC 7975 | pMQ30 $\Delta mapA$ | This study |
| SMC 7907 | pMQ30 MapA-HA | This study |
| SMC 8703 | pMQ30 $\Delta mapE$ | This study |
| SMC 8637 | pMQ30 $P_{mapA} :: P_{mapA}$ | This study |
| SMC 8635 | pMQ56 mTn7 *plapA*-GFP, *pmapA*-mCherry | This study |
| SMC 6174 | pMQ72 *gcbC*-R366E | (63) |

### Biofilm assay

*P. fluorescens* strains were struck out on LB plates overnight. Single colonies were picked and grown overnight in liquid LB medium with shaking. 1.5µl of overnight culture was added to 100µl of growth medium in a 96-well U-bottom polystyrene plate (Costar). Plates were covered and placed in a humidified microchamber at 30°C for either 6 or 16 hours as indicated, at which time the liquid in the wells was discarded and the wells were stained with 125µl of 0.1% (w/v) crystal violet (CV) at room temperature for 20 min and then rinsed 2 times with water. Wells were allowed to dry and the stain was dissolved using 150µl of a solution of water, methanol, and acetic acid (45:45:10 ratio by volume) for 10 min at room temperature. A 100µl volume of the solubilized CV solution was transferred to a flat-bottom 96-well plate, and the OD was recorded at 550 nm.

### Construction of HA-tagged MapA

The 3XHA tag was amplified by PCR from *P. fluorescens* Pf0-1 in which LapA had been HA-tagged. This tag was inserted into the genomic copy of the Pfl01_1463 (*mapA*) gene between the codon encoding H2601 and G2602 by allelic exchange, as previously described (86).

### Assessment of surface localization of adhesins by dot blot

WT *P. fluorescens* Pf0-1 and mutants were grown with agitation in LB at 30°C for ∼16 hours, then 100µl of this inoculum was added to 25mL of KA medium. This subculture was incubated with agitation at 30°C for 24 hours. After incubation, the 25mL cultures were centrifuged at 5000 rpm for 5 minutes, and cell pellets were resuspended in 1mL of fresh medium. All cultures were washed once with fresh medium and normalized such that the OD_600_ of each culture was the same. 5ul of washed, normalized culture was spotted onto a nitrocellulose membrane and allowed to dry. The membrane was then incubated in blocking buffer for 1 hour at 25°C (TBST with 3% BSA; Sigma) before being probed for LapA or MapA containing an HA tag with an anti-HA antibody (Biolegend) at a 1:2000 dilution in TBST with 3% BSA and 0.02% sodium azide overnight at 4°C. Excess antibody was washed away with 3 x TBST rinses before the membrane was incubated with anti-mouse secondary antibody (Bio-rad) conjugated to horseradish peroxidase at a 1:15000 dilution in TBST at 25°C for 1 hour. The membrane was then washed 3 times in TBST and once in TBS (Bio-rad) before being exposed to 2ml of Western Lightning Plus-ECL, Enhanced Chemiluminescence Substrate (Perkin-Elmer) The membrane was then exposed to Biomax XAR Film (Carestream) and the film was processed on a Kodak X-OMAT 2000 Film Processor. The processed films were scanned in grayscale, and the dots were quantified in ImageJ (NIH) by reversing the pixel values, defining a region of interest (ROI), and then using that defined object to measure the mean gray value of each spot and the background of the film, which was defined as an area of the film where no spot was present. The background measurement was subtracted from the mean pixel density of each spot.

### Nanostring transcriptional measurement and analysis

For each biological replicate, 11 spots of 5µl each from each overnight LB liquid culture of the wild-type and mutant strains were spotted onto 1.5% agar plates containing K10T-1 medium. Cells were grown at 30°C for 6 h, scraped from the plate, and resuspended in Tris-EDTA (TE) buffer. All of the 11 spots were pooled into one biological replicate to obtain sufficient RNA for the analysis, and this experiment was performed on three different days to attain three biological replicates. The cells were processed using a RNeasy extraction kit (Qiagen), and 75ng of each RNA sample was added to a Nanostring nCounter kit, and the protocol provided by the manufacturer (Nanostring) was followed without modification. Raw reads from the Nanostring system were normalized as a fraction of the total number of reads measured for targeted genes, expressed as reads per 1,000 transcripts as described previously (87).

### Microfluidic devices

Microfluidic devices were constructed as previously described (88). Specifically, the devices were constructed by bonding polydimethylsiloxane (PDMS) castings to size #1.5 36 mm x 60 mm cover glass (ThermoFisher, Waltham MA) (89, 90). Bacterial strains were grown in 5ml LB with 15µg/µl gentamycin overnight at 30 °C with agitation. 300ul of overnight culture was pelleted and the supernatant removed before being resuspended in 1ml the medium described above. Strains were inoculated into channels of the microfluidic devices and allowed to colonize for 1 hour at room temperature 21-24°C before providing constant flow of 0.5ul/minute of FCM. Medium flow was achieved using syringe pumps (Pico Plus Elite, Harvard Apparatus) and 5ml syringes (27-guage needle) fitted with #30 Cole palmer PTFE tubing (ID = 0.3mm). Tubing was inserted into holes bored in the PDMS with a catheter punch driver.

### Imaging

Biofilms were imaged using an Andor W1 Spinning Disk Confocal, Andor Zyla camera, and Nikon Ti microscope. A 488nm laser was used to excite GFP, while a 560nm laser line was used to excite mRuby. Images were captured using and Nikon NIS software.

### LapD and LapG homolog prediction and LapA-like protein identification

The approach to identifying LapD and LapG encoding organisms and LapA-like proteins was modified from a previously described approach (29). The code is posted at https://github.com/GeiselBiofilm/Collins-MapA. Briefly, LapG and LapD homologs were defined as ORF-coding proteins with the pfam06035 and pfam16448 domains, respectively. The NCBI Conserved Domains Database (CDD) was utilized to generate a list of LapG- and LapD-encoding bacteria, and the programming language R (91) was used to determine the intersecting LapD- and LapG-encoding bacteria by text matching their species name identifiers. The protein annotations of these genomes were downloaded from the NCBI genome database, and each annotated locus with a size of greater than 1000 amino acids was interrogated for the presence of a LapG cleavage site within amino acids 70 to 150 ([T/A/P]AA[G/V]) and at least one RTX motif (DX[L/I]X4GXDX[L/I]XGGX3D).

### Phylogenetic tree construction

Representatives of each *Pseudomonas* species in the GenBank database were selected based on genome quality (i.e. fully assembled genome, or highest n50 if genome in contigs or scaffolds). In the case of species for which multiple strains with high quality genomes were available, common lab strain representatives were chosen. A list of the strains used in the construction of this tree is given in supplemental file 2. The genome assembly files were retrieved from the GenBank ftp server and used to construct a phylogenetic tree according to a previously described method as follows (72, 73). Orthologs of the 16S rRNA, *gyrB, rpoB*, and *rpoD* genes were identified by BLASTn version 2.6.0 (using the setting: -gapopen 2) using the sequences of *P. fluorescens* Pf0-1 genes as queries (Pfl01_R12 : 16S rRNA, Pfl01_0004: *gyrB*, Pfl01_5086: *rpoB*, Pfl01_5149: *rpoD*). Any species for which no BLAST matches could be found for any of the query genes were removed. Multiple sequence alignment was conducted using MUSCLE version 3.8.31 (74) for each gene, and alignments were concatenated. Concatenated aligned sequences were analyzed using PAUP*4.0a (build 167)(92). Distance was calculated using a Jukes-Cantor model and a tree was constructed using neighbour-joining.

## Acknowledgement

We thank Carey Nadell and his laboratory for the assistance with microfluidic chambers and imaging. The work was supported by NIH grant R01 GM123609 to G.A.O. We would also like to acknowledge the Molecular Biology Core who receives funding from the Norris Cotton Cancer Center Core grant from the NIH (P30-CA023108).

